# Effects of exogenous β-glucanase on ileal digesta soluble β-glucan molecular weight, digestive tract characteristics, and performance of coccidiosis challenged broiler chickens fed hulless barley-based diets with and without medication

**DOI:** 10.1101/2020.07.06.189258

**Authors:** Namalika D. Karunaratne, Rex W. Newkirk, Nancy P. Ames, Andrew. G. Van Kessel, Michael R. Bedford, Henry L. Classen

## Abstract

Limited use of medication in poultry feed led to the investigation of exogenous enzymes as antibiotic alternatives for controlling enteric disease. The objective of this study was to evaluate the effects of diet β-glucanase (BGase) and medication on β-glucan depolymerization, digestive tract characteristics, and performance of broilers. Broilers were fed hulless barley (HB) based diets with BGase (Econase GT 200P from AB Vista; 0 and 0.1%) and medication (Bacitracin and Salinomycin Na; with and without) arranged as a 2 × 2 factorial. In Experiment 1, 160 broilers were housed in cages from d 0 to 28. Each treatment was assigned to 10 cages. In Experiment 2, broilers (2376) were housed in floor pens and challenged with a coccidiosis vaccine on d 5. Each treatment was assigned to one floor pen in each of nine rooms. In Experiment 1, the soluble β-glucan weight average molecular weight (Mw) in the ileal digesta was lower with medication in the 0% BGase treatments. Peak molecular weight (Mp) and Mw were lower with BGase regardless of medication. The maximum molecular weight for the smallest 10% β-glucan (MW-10%) was lower with BGase. In Experiment 2, Mp was lower with medication in 0% BGase treatments. Beta-glucanase resulted in lower Mp regardless of medication, and the degree of response was lower with medication. The MW-10% was lower with BGase despite antibiotic addition. Body weight gain (BWG) and feed efficiency were higher with medication regardless of BGase use through-out the trial (except d 11-22 feed efficiency). Beta-glucanase resulted in higher BWG after d 11, and lower and higher feed efficiency before and after d 11, respectively, in unmedicated treatments. In conclusion, BGase and medication caused the depolymerization of soluble ileal β-glucan. Beta-glucanase appeared as a partial replacement for diet medication to increase coccidiosis challenged broiler performance.

## Introduction

Antibiotics have been used in poultry feed at sub-therapeutic doses for decades to improve growth and feed efficiency, and to prevent enteric infections [1]. However, the prolonged and indiscriminate use of antimicrobials in animal production is likely to cause the development of antibiotic resistance in pathogenic bacteria, and its’ effect on animal and human health risk has led to reduced use of in-feed antibiotics in the poultry industry [2, 3]. Therefore, the investigation of alternatives to antibiotics is a primary focus to control infectious enteric diseases and promote growth and gut health in poultry [4, 5]. Potential alternatives to antibiotics that have been studied include probiotics, prebiotics, organic acids, essential oils and feed enzymes [6, 7].

Prebiotics are non-digestible feed ingredients that beneficially affect the host by selectively stimulating the growth and function of beneficial microbiota in the digestive tract [8]. The most commonly available prebiotics are oligosaccharides from various sources, and small molecular weight polysaccharides derived from cereal grains. Studies in the literature have focused on molecules such as fructo-oligosaccharides, mannose-oligosaccharides, xylo-oligosaccharides and arabinoxylo-oligosaccharides in terms of improving poultry digestive tract health and production performance, and modulating intestinal microbiota, epithelial integrity, and immune function in poultry. Dietary mannan-oligosaccharides have been shown to increase morphological development of the digestive tract and colonization of beneficial bacteria while reducing pathogenic bacteria in chickens [9, 10]. Fructo-oligosaccharides have also demonstrated beneficial effects on broiler chickens in terms of intestinal epithelial morphology [11, 12], digestive tract microbiota [13, 14], and bird immune response [15, 16]. Dietary inclusion of arabinoxylo-oligosaccharides/ xylo-oligosaccharides affects gastro-intestinal microbial populations of chickens by increasing beneficial bacteria, including Bifidobacteria, Lactobacilli and *Clostridium* cluster XIV [17, 18], and reducing *Salmonella* colonization in the caeca and translocation to the spleen [19]. In addition, exogenous xylanase in wheat-based diets increased the number of gastro-intestinal beneficial bacteria, including lactic acid bacteria, while reducing pathogenic bacteria in broiler chickens [20, 21], probably by decreasing the molecular weight of soluble arabinoxylan derived from the wheat. Arabinoxylan has been extensively studied concerning its ability to act as a prebiotic since arabinoxylan is found in the cell walls of the most common cereals used in poultry feed (wheat and corn) and prebiotic oligosaccharides are presumed to be formed on use of a xylanase. However, research is limited regarding cereal β-glucan since it predominates in barley and oats, which are less commonly found in poultry feed. Therefore, it is relevant to investigate the effects of low molecular weight barley β-glucan produced by supplementing exogenous β-glucanase (BGase) in broilers fed barley-based diets.

Hulless barley (HB) contains a higher level of β-glucan compared to conventional barley due to the removal of the hull during processing [22, 23]. Further, many HB cultivars are developed for the human food industry, and as a result, are selected for high β-glucan content [24]. Dietary enzymes such as endo-β-glucanase depolymerize larger molecular weight β-glucan producing lower molecular weight compounds, which are fermentable in the distal digestive tract [25]. A consequence of fermentation is the production of short-chain fatty acids (SCFA), which are thought to improve digestive tract morphology and physiology and stimulate the establishment of beneficial bacterial populations, while at the same time reducing colonization by pathogens [26, 27]. However, the effects of exogenous BGase on microbial fermentation and digestive tract physiology and morphology are less-well studied, and the results have been inconsistent in previous research. Therefore, investigating the mechanism of action of diet BGase on HB β-glucan might contribute to the understanding of the enzyme effect on the digestive tract characteristics of chickens.

The mechanisms of action of feed medication are not fully understood, although antibiotics have been successfully used to promote growth and feed efficiency and improve bird health [16, 28]. The primary mechanism is generally accepted to be a positive modulation of the diversity and relative abundance of bacteria in the digestive tract microbial community, and thereby the control of enteric disease and stimulation of immune function in broiler chickens [29–31]. However, other beneficial mechanisms are also possible. Investigating the interaction between medication and enzyme use in high fibre diets offers the potential to add knowledge on medication mechanisms of action and to study the effectiveness of enzymes in reducing the adverse effects of enteric disease. The effects of exogenous BGase and diet medication on broiler performance and digestive tract characteristics could depend on the age of the birds due to the distinct maturity of the digestive tract, including the development of gut microbiota, and housing conditions that affect the level of exposure to pathogenic organisms. Therefore, the current study utilized the same experimental design and treatments in two different environments. Experiment 1 was completed in battery cages and a low disease challenge environment, while Experiment 2 was completed in litter floor pens using broilers challenged with a vaccine against coccidiosis at 5 d of age. The rationale for these experiments was to determine if treatments produce the same effects in the prescribed settings.

The objective of the current study was to investigate the effects of exogenous BGase and medication on ileal digesta soluble β-glucan molecular weight distribution, digestive tract characteristics, and production performance of broiler chickens fed an HB-based diet under different housing environments and disease conditions. It was hypothesized that exogenous BGase would depolymerize high molecular weight β-glucan, resulting in increased fermentation in the distal digestive tract and beneficial effects on the digestive tract morphology and physiology. This should result in improved production performance of broiler chickens and thereby reduce the requirement for medication in broilers fed HB-diets. Further, a higher response to exogenous BGase and a greater reduction of the necessity of diet medication would be expected from the broiler chickens from Experiment 2 (coccidiosis-challenged) compared to Experiment 1 due to increased environmental pressures.

## Materials and methods

The experimental procedure was approved by the Animal Research Ethics Board of the University of Saskatchewan and conducted according to the Canadian Council on Animal Care guidelines for humane animal use [32, 33].

### Birds and housing

#### Experiment 1

A total of 160 broiler chickens (Ross × Ross 308) obtained from a commercial hatchery were housed in battery cages. The dimensions of the cages were 51 cm in length, 51 cm in width and 46 cm in height. The grid size of the wire mesh floor of each cage was 2.54 × 2.54 but was covered by a 1.27 × 1.27 cm mesh until d 7. There were two levels of battery cages that were in two rows with back to back cages. The starting room temperature was 32°C, and it was gradually decreased by 2.8°C per week. The minimum light intensity was 25 lux during the experimental period, and the day length was 23 h (d 0-7) and 18 h (d 8-28). Birds were given feed and water *ad-libitum* throughout the experiment. Each cage had a front-mounted feed trough (51 cm in length) and two height-adjustable nipple drinkers. Extra feed and water were supplied to the birds from d 0 to 5 using supplementary chick feeders (50 cm long, plastic) and ice cube trays (16 cells), respectively. There were 10 cage replications per treatment and four birds per cage. Treatments were randomly assigned to the battery cages.

#### Experiment 2

A total of 2376 one d old male and female (Ross × Ross 308) broiler chickens were obtained from a commercial hatchery and randomly placed in 36 floor pens (2.3 m × 2.0 m) in nine environmentally controlled rooms. Each room contained four pens randomly assigned to the four treatments; each treatment was replicated nine times. Each pen (66 birds per pen) contained a tube feeder (pan diameter - 36 cm from 0 to 25 d and 43 cm after that) and a height-adjustable nipple drinker (six Lubing nipples). Additional feed and water were supplied to each pen using a cardboard egg tray and an ice cube tray, respectively, for the first week. Straw was placed in each room at a thickness of 7.5-10 cm. The room temperature was 33°C at the beginning of the experiment and was gradually reduced to 21°C by d 25. Day length was gradually reduced from 23 h at d 0 to 17 h at d 12, and the light intensity was set to 20 lux at the start of the experiment and gradually decreased to 10 lux by d 10. Birds were given feed and water *ad-libitum* throughout the experiment.

### Experimental diets

The dietary treatments were arranged according to a 2 × 2 factorial arrangement (BGase and medication) in both experiments. Beta-glucanase (Econase GT 200 P from ABVista, Wiltshire, UK) levels were 0 and 0.1% (the β-glucanase activity of 0 and 200,000 BU/kg, respectively), and diets were fed without or with medication (Bacitracin (Zoetis Canada Inc., Kirkland, QC, Canada) at 4.4 mg/kg and Salinomycin Sodium (Phibro Animal Health Corporation, Teaneck, NJ) at 25 mg/kg). Diets were based on 60% hulless barley (CDC Fibar) and were formulated to meet or exceed Ross 308 broiler nutrition specifications [34]. The ingredients and calculated nutrient levels are shown in Table 1. Diets were fed in crumble form in Experiment 1. In Experiment 2, starter diets (d 0-11) were fed in crumble form, and grower diets (d 11-33) were given initially in crumble form, and then switched to a pellet form. The pelleting temperature was controlled between 70-75°C to prevent high temperature-induced BGase inactivation during feed processing. Measured β-glucanase activity in diets approached the estimated values in both experiments, thereby confirming β-glucanase was added correctly, and that activity was not lost during feed processing. Xylanase activity was non-detectable in experimental diets.

**Table 1.** Ingredients and calculated nutrient levels (%) of Experimental diets

| **Ingredient** | **Experiment 1** | **Experiment 2** |  |
| --- | --- | --- | --- |
|  |  | **Starter** | **Grower** |
| Hulless barley | 60.00 | 59.09 | 60.00 |
| Wheat | 4.46 | 0.00 | 4.55 |
| Soybean meal | 26.93 | 32.97 | 26.99 |
| Canola oil | 4.07 | 3.29 | 4.13 |
| Monocalcium phosphate | 1.20 | 1.40 | 1.20 |
| Limestone | 1.52 | 1.64 | 1.52 |
| Sodium chloride | 0.38 | 0.43 | 0.38 |
| Vitamin-mineral broiler premix<sup>1</sup> | 0.50 | 0.50 | 0.50 |
| Choline chloride | 0.10 | 0.10 | 0.10 |
| DL-Methionine | 0.27 | 0.30 | 0.27 |
| L-Threonine | 0.05 | 0.07 | 0.05 |
| L-Lysine HCl | 0.22 | 0.21 | 0.22 |
| **Nutrient, calculated** |  |  |  |
| AME (kcal/kg) | 3100 | 3000 | 3100 |
| Crude protein | 21.24 | 23.46 | 21.24 |
| Crude fat | 5.57 | 4.74 | 5.57 |
| Calcium | 0.87 | 0.96 | 0.87 |
| Chloride | 0.36 | 0.38 | 0.36 |
| Non-phytate phosphorous | 0.44 | 0.48 | 0.44 |
| Potassium | 0.83 | 0.92 | 0.83 |
| Sodium | 0.18 | 0.20 | 0.18 |
| Digestible arginine | 1.35 | 1.50 | 1.35 |
| Digestible isoleucine | 0.81 | 0.90 | 0.81 |
| Digestible leucine | 1.47 | 1.61 | 1.47 |
| Digestible lysine | 1.15 | 1.28 | 1.15 |
| Digestible methionine | 0.54 | 0.60 | 0.54 |
| Digestible methionine and cysteine | 0.87 | 0.95 | 0.87 |
| Digestible threonine | 0.77 | 0.86 | 0.77 |
| Digestible tryptophan | 0.24 | 0.27 | 0.24 |
| Digestible valine | 0.87 | 0.96 | 0.87 |

<sup>1</sup>Vitamin-mineral premix provided the following per kilogram of complete diet: vitamin A, 11,000 IU; vitamin D<sub>3</sub>, 2,200 IU; vitamin E, 30 IU; menadione, 2 mg; thiamine, 1.5 mg; riboflavin, 6 mg; pyridoxine, 4 mg; vitamin B<sub>12</sub>, 0.02 mg; niacin, 60 mg; pantothenic acid, 10 mg; folic acid, 0.6 mg; biotin 0.15 mg; copper, 10 mg; iron, 80 mg; manganese 80 mg; iodine, 0.8 mg; zinc, 80 mg; selenium, 0.3 mg; calcium carbonate 500 mg; ethoxyquin 0.63 mg; wheat middlings 3773 mg.

### Coccidiosis challenge

In Experiment 2, all the birds were challenged with the Coccivac B-52 live vaccine (Merck Animal Health; 1.3× recommended dose). The vaccination was completed at d 5 to facilitate uniform intake of coccidian oocysts by the birds. The vaccine contains oocysts of *Eimeria acervulina*, *E. mivatis*, *E. maxima* and *E. tenella*. The vaccine was sprayed on feed located in a cardboard egg tray and into water placed in an ice cube tray. A 30 cm wide Kraft brown paper strip (Model S-8511S, ULINE Canada, Milton, Ontario, Canada) was placed under the full length of the nipple drinker line in each pen before vaccination to facilitate oocyst ingestion by the birds. In addition, 60% relative humidity was maintained in the rooms, using humidifiers and water application, to facilitate oocyst cycling. Feeders and drinkers were raised in each pen before vaccination and were put-down once the vaccine containing feed and water was consumed by the birds.

### Performance data collection

Body weight and feed intake (FI) were measured on a cage basis at d 7, 14, 21 and 28 in Experiment 1. In Experiment 2, body weight and FI were measured on a pen basis at d 11, 22 and 32. Mortality was recorded daily, and dead birds were sent to Prairie Diagnostic Services for necropsy.

### Sample collection

In Experiment 1, all birds were euthanized on d 28, whereas in Experiment 2, a total of four birds per pen were euthanized at two collection points (d 11 and 33). Birds were euthanized by administering T-61 (Embutramide, Mebezonium iodide and tetracaine; Merck animal health, Kirkland, Quebec, Canada) into the brachial vein. Birds were weighed individually. Two birds were used for pH measurement and to collect samples for SCFA analysis and histology (in Experiment 2 only) at each collection. *In-situ* pH of the crop, gizzard, duodenum, jejunum, ileum, caeca and colon contents was measured using a Beckman Coulter 34 pH meter (Model PHI 34, Beckman Instruments, Fullerton, CA). Two 1 cm samples of mid-ileum were sectioned, placed in 10% neutral buffered formalin, and stored at room temperature until histo-morphology evaluation. Total ileal and caecal contents were collected into plastic centrifuge tubes and stored at −20°C for the analysis of SCFA. Two birds were used to collect digestive tract size, content, and organ data at each collection in both trials. The digestive tract was detached from the bird carcass and then sectioned into the crop, proventriculus, gizzard, duodenum, jejunum, ileum, caeca and colon; the liver, spleen and pancreas were removed and weighed. Full and empty weights of all sections and the length of each intestinal section were recorded. The content weight of each section was determined by subtracting empty weight from the full weight. Relative tissue weights and lengths were calculated based on individual bird weight. Total ileal contents were collected into plastic snap-cap vials (pooled from all the birds in a cage in Experiment 1; one bird per pen in Experiment 2) and centrifuged for 5 min at 17013 × g using a Beckman microfuge (Model E 348720, Beckman Instruments, INC, Palo Alto, CA). Then the viscosity of ileal supernatant was measured using a Brookfield cone-plate digital viscometer (Model LVDV-Ⅲ, Brookfield Engineering Labs, INC, Stoughton, MA 02072), which was maintained at 40⁰C (40 rpm; shear rate 300 s^-1^). The rest of the ileal supernatant was stored at - 80°C for β-glucan molecular weight distribution analysis.

### Nutritional analysis

The ingredients (HB and wheat) were ground using a Retsch laboratory mill (Retsch ZM 200, Germany) and analyzed for total starch, CP, fat, ash, moisture and fibre following AOAC, AACC and ICC standard methods [35–37]. Ingredients were analyzed for total starch using the AOAC method 996.11 and the AACC method 76-13.01 using a Megazyme kit (Total starch assay procedure, Amyloglucosidase/α-amylase method, Megazyme International Ireland Ltd., Bray Business Park, Bray, Co. Wicklow, Ireland). Nitrogen was analyzed using a Leco nitrogen analyzer (Model Leco-FP-528L, Leco Corporation, St. Joseph, MA, USA), and 6.25 was the N to CP conversion factor. Ether extraction was completed using Goldfish Extraction Apparatus (Labconco model 35001; Labconco, Kansas, MO, USA) following the AOAC method 920.39 to determine fat content. Ash content was analyzed according to the AOAC method 942.05 using a muffle oven (Model Lindberg/Blue BF51842C, Asheville, NC 28804, USA). Moisture was analyzed using the AOAC method 930.15. The analysis of insoluble dietary fibre (IDF) and soluble dietary fibre (SDF) was completed using a Megazyme kit (Total dietary fibre assay procedure, Megazyme International Ireland Ltd., Bray Business Park, Bray, Co. Wicklow, Ireland) according to the AOAC method 991.43 and the AACC method 32-07.01. Total dietary fibre (TDF) was obtained by adding IDF and SDF. Beta-glucan was analyzed using a Megazyme analysis kit (Mixed-linkage beta-glucan assay procedure/McCleary method, Megazyme International Ireland Ltd., Bray Business Park, Bray, Co. Wicklow, Ireland) according to the AOAC Method 995.16, AACC Method 32-23, and ICC Standard Method No. 168. In addition, diets were analyzed for β-glucanase (EC 3.2.1.6) and xylanase activity (EC 3.2.1.8) according to the AB Vista methods of ESC Standard Analytical Methods SAM042-01 and SAM038, respectively (ABVista, Wiltshire, UK).

### Beta-glucan molecular weight distribution

Ileal supernatant samples were boiled for 15 min and centrifuged at 17,013 × g for 10 min using a Beckman microfuge (Model E348720, Beckmann instruments, INC, Palo Alto, CA). The sample was then analyzed for β-glucan molecular weight using size exclusion chromatography and calcofluor post-column derivatization [38]. The two columns used for HPLC were Shodex OHpak SB-806M with OHpak SB-G column guard and a Waters Ultrahydrogel linear column. The mobile phase was 0.1M Tris buffer (pH=8). Molar mass distribution curves were used to obtain β-glucan Mp, weight average molecular weight (Mw), and the maximum molecular weight for the smallest 10% β-glucan molecules (MW-10%) of each sample. Peak molecular weight is the molecular weight of the highest β-glucan fraction. Weight average molecular weight is the average of the molecular weights of all β-glucan molecules (considering the weight fraction of each type of molecule).

### Short chain fatty acids analysis

Short chain fatty acids were analyzed in triplicate according to the procedure described by [39] with minor changes. The internal standard for the analysis was made up of 20 ml of 25% phosphoric acid, 300 µl of isocaproic acid, and deionized water. Three hundred microliters of acetic acid, 200 µl of propionic acid, 100 µl of butyric acid, and 50 µl of isobutyric, isovaleric, valeric, caproic and lactic acids were used to make the standard solution. The digesta was thawed and mixed with 25% phosphoric acid at 1:1 and kept at room temperature for 10 min with occasional shaking. It was then centrifuged at 12,500 × g for 10 min. The supernatant (1 ml) was mixed with 1 ml of the internal standard and centrifuged at 12,500 × g for 10 min. It was filtered using a 0.45-micron nylon filter, and the filtrate was placed in a GC autosampler vial and injected into a Zebron Capillary Gas Chromatography column (length 30m, internal diameter 0.25 mm, film thickness 0.25 µm; (Zebron^TM^ZB-FFAP, Phenomenex, Torrance, CA). The SCFA analysis was completed using the Thermo Scientific Gas Chromatography system (Model Trace 1310, Milan, Italy).

### Histomorphology of gastro-intestinal wall

In Experiment 2, ileal tissue samples were cut into two longitudinal sections and embedded in paraffin. Two slides were made from each sample to obtain ileal morphology measurements (Hematoxylin and Eosin stain) and goblet cell (GC) categorization (Alcian Blue/ Periodic Acid-Schiff stain). An Optika B-290TB digital microscope (Bergamo, Italy) was used to observe slides, and an HDCE-X3 digital camera with Optika Vision Lite software was used to capture the images. Well-oriented 8-10 villi and crypts per section were used to measure villi length, width, and crypt depth. Villi length was considered as the length from the tip of a villus to the villus-crypt junction. The villi width was measured at the middle of the villus height. The depth of the invagination between adjacent villi was considered as the crypt depth. Goblet cells were counted around the perimeter of 8-10 well-oriented villi per section, and the three categories of GC were identified, acidic mucin-producing GC (stained in blue), neutral mucin-producing GC (stained in magenta) and mixed mucin-producing GC (stained in purple) [40].

### Statistical analysis

Data were analyzed using the Proc Mixed model of SAS 9.4 [41]. Both experiments were randomized complete block designs, and the battery cage level and room were considered as blocks for Experiments 1 and 2, respectively. Treatments were replicated 10 times in Experiment 1 (battery cages equally distributed in two levels), and nine times in Experiment 2 (one pen in nine different rooms). Differences were considered significant when *P* ≤ 0.05. Data were checked for normality and analyzed using 2-way ANOVA. Tukey-Kramer test was used to detect significant differences between means.

## Results

### Ingredient nutrient composition

In Experiment 1, TDF, IDF, SDF and total β-glucan in HB were 29.0, 19.6, 9.6 and 8.70%, respectively, and the same parameters were 15.2, 13.7, 1.6 and 0.68%, respectively for wheat. The content of total starch, CP, fat and ash were measured as 49.7, 16.2, 2.4 and 2.4%, respectively, in HB, and as 64.1, 15.0, 1.2 and 1.9% in wheat. In Experiment 2, TDF, IDF, SDF and total β-glucan were 26.7, 18.9, 7.8 and 8.70% (HB); 14.4, 12.4, 2.0 and 0.64% (wheat), respectively. In addition, total starch, CP, fat and ash were determined to be 53.7, 16.2, 2.8 and 2.4% in HB, and as 62.8, 14.9, 1.2 and 1.7% in wheat, respectively.

### Beta-glucan molecular weight distribution

In Experiment 1, both Mp and Mw were affected by the interaction between main effects; values were lower with enzyme use regardless of diet medication, but the degree of response was less in medicated diets. In addition, Mw was lower with the use of medication when the birds were given diets without BGase. The MW-10% values were unaffected by medication but were lower with 0.1% compared to 0% BGase.

In Experiment 2, interactions were found for all molecular weight criteria at both ages (11 and 33 d) except for Mw at 11 d, which was also unaffected by medication or BGase. Values for Mp and Mw-10% followed a similar trend, with enzyme consistently reducing values, but with the degree of response less in medicated diets. In the absence of the enzyme, medication reduced Mp at both ages and MW-10% on d 33. The interaction for Mw at 33 d was due to enzyme decreasing and increasing Mw for nonmedicated and medicated diets, respectively.

Figures 1A and 1B compare the β-glucan molecular weight distribution of ileal digesta from 11 d broilers fed diets without medication, and without and with BGase, respectively, in Experiment 2. Beta-glucanase increased the proportion of low molecular weight β-glucan, as shown by curve placement relative to the blue line at x-axis point 1e^4^. Diet medication also increased the proportion of low molecular weight β-glucan in comparison to the nonmedicated diet, and this is contrasted in Figs 1A and 1C.

**Figure 1.**
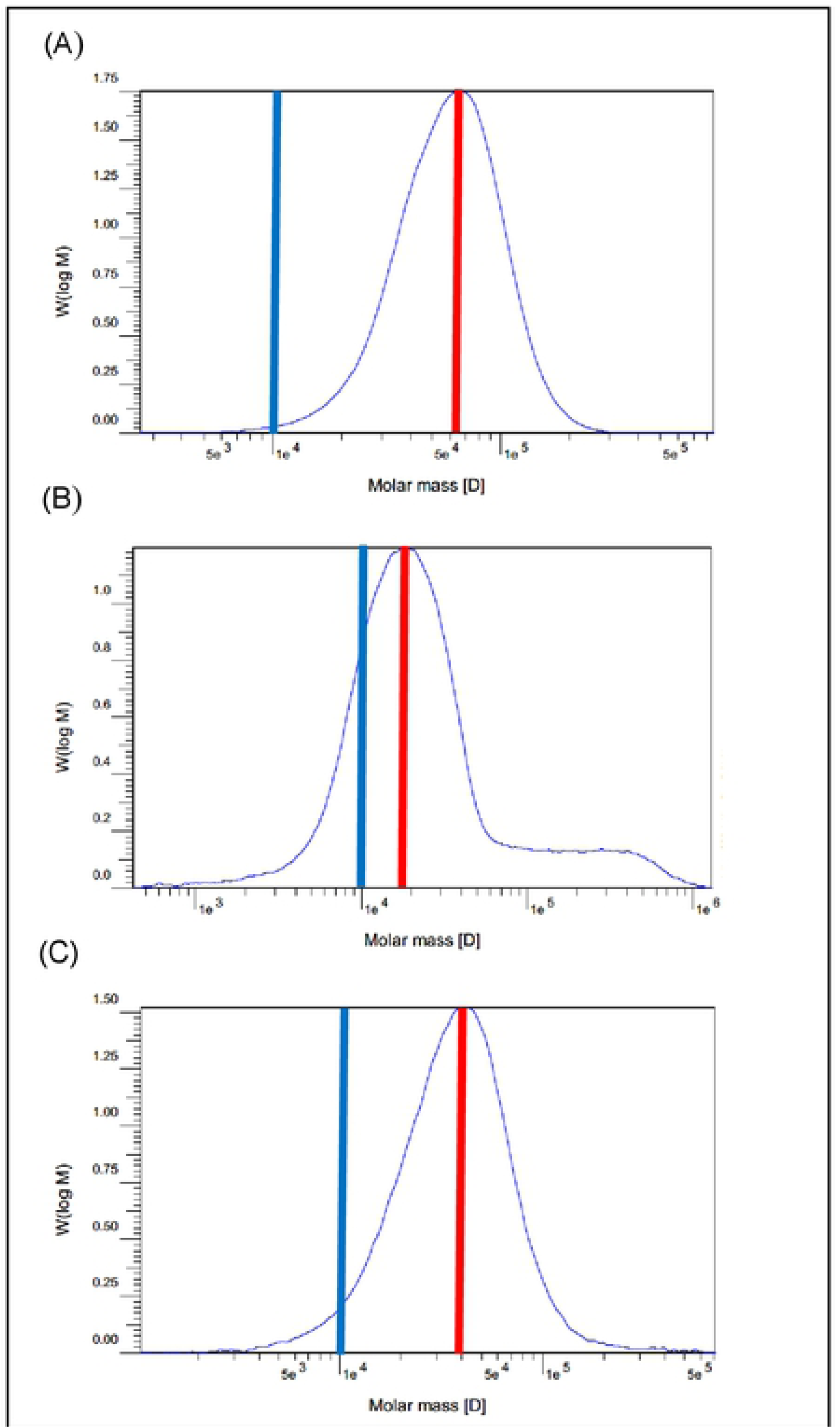
Beta-glucan molecular weight distribution in soluble ileal digesta from 11 d broilers fed 60% hulless barley diets in Experiment 2. Blue lines denote point 1e^4^ on the x-axis and red lines indicate the Mp of the distribution curve. (A) Without medication, 0% β-glucanase (B) Without medication, 0.1% β-glucanase (C) With medication, 0% β-glucanase

### Viscosity

Ileal digesta viscosity was not affected by medication in Experiment 1 but was reduced with the use of BGase. In Experiment 2 at 11 d, an interaction was found between medication and BGase; BGase reduced viscosity without dietary medication. In the interaction, the highest viscosity was noted for the treatment without medication or BGase, and the lowest was the treatments with BGase; treatment with medication and without BGase was intermediate. At d 33 in Experiment 2, BGase decreased viscosity, but there was no medication effect.

### Short chain fatty acids and gastro-intestinal pH

Ileal digesta SCFA levels and molar percentages were not affected by dietary treatments in Experiment 1, except for caproic acid concentration, where values were lower with BGase supplementation (Table 4). Similarly, caecal digesta SCFA concentrations and molar percentages were also not affected by treatment (Table 5). Noteworthy, the interaction between medication and BGase tended to be significant (*P* = 0.06-0.09) for the concentrations of total and individual SCFA. In all cases, levels tended to decrease with enzyme use in the non-medicated diets and increase with enzyme use in the medicated diets.

**Table 2.** Effects of diet medication and β-glucanase on β-glucan molecular weight in ileal content of broiler chickens

| Medication | β-glucanase (%) | Molecular weight (g/mol) | | | | | | | | |
| --- | --- | --- | --- | --- | --- | --- | --- | --- | --- | --- |
|  |  | Experiment 1 |  |  | Experiment 2 |  |  |  |  |  |
|  |  | d 28 |  |  | d 11 |  |  | d 33 |  |  |
|  |  | Mp<sup>1</sup> | Mw | MW-10% | Mp | Mw | MW-10% | Mp | Mw | MW-10% |
| without | 0 | 19799<sup>a</sup> | 36199<sup>a</sup> | 6096 | 78293<sup>a</sup> | 80971 | 33322<sup>a</sup> | 65176<sup>a</sup> | 69508<sup>a</sup> | 29025<sup>a</sup> |
|  | 0.1 | 7793<sup>b</sup> | 8434<sup>c</sup> | 1955 | 24568<sup>c</sup> | 63835 | 7250<sup>b</sup> | 16985<sup>c</sup> | 48316<sup>b</sup> | 7074<sup>c</sup> |
| with | 0 | 16824<sup>a</sup> | 19119<sup>b</sup> | 5326 | 54475<sup>b</sup> | 59002 | 26065<sup>a</sup> | 40595<sup>b</sup> | 49017<sup>b</sup> | 13586<sup>b</sup> |
|  | 0.1 | 10401<sup>b</sup> | 9929<sup>c</sup> | 2201 | 27677<sup>c</sup> | 61898 | 10586<sup>b</sup> | 22144<sup>c</sup> | 60641<sup>a</sup> | 8157<sup>c</sup> |
| SEM<sup>2</sup> |  | 1148.1 | 2513.9 | 509.2 | 5982.7 | 3537.4 | 2717.0 | 4481.7 | 2258.9 | 1890.1 |
| Main effects |  |  |  |  |  |  |  |  |  |  |
| <u>Medication</u> |  |  |  |  |  |  |  |  |  |  |
| without |  | 13796 | 22317 | 4025 | 51431 | 72403 | 20286 | 41080 | 58912 | 18049 |
| with |  | 13612 | 14524 | 3763 | 41076 | 60450 | 18325 | 31370 | 54829 | 10871 |
| <u>β-glucanase (%)</u> |  |  |  |  |  |  |  |  |  |  |
| 0 |  | 18311 | 27659 | 5711<sup>a</sup> | 66384 | 69986 | 29694 | 52885 | 59263 | 21305 |
| 0.1 |  | 9096 | 9181 | 2078<sup>b</sup> | 26122 | 62867 | 8918 | 19565 | 54479 | 7615 |
| <u>Probability</u> |  |  |  |  |  |  |  |  |  |  |
| Medication |  | 0.86 | 0.001 | 0.70 | 0.08 | 0.06 | 0.39 | 0.04 | 0.16 | <.0001 |
| β-glucanase | | <.0001 | <.0001 | <.0001 | <.0001 | 0.21 | <.0001 | <.0001 | 0.10 | <.0001 |
| Medication × β-glucanase | | 0.01 | 0.0004 | 0.45 | 0.03 | 0.09 | 0.03 | 0.004 | <.0001 | <.0001 |

<sup>a-c</sup>Means within a main effect or interaction not sharing a common superscript are significantly different ( $P \leq 0.05$ ).
<sup>1</sup>Mp - peak molecular weight; Mw - weight average molecular weight; MW-10% - The maximum molecular weight for the smallest 10% molecules.
<sup>2</sup>SEM - pooled standard error of mean (d 28, n=6 cages per treatment; d 11 and 33, n=6 birds per treatment).

**Table 3.** Effects of diet medication and β-glucanase on the ileal soluble digesta viscosity of broiler chickens

| Medication | β-glucanase (%) | Viscosity (cP) | | |
| --- | --- | --- | --- | --- |
|  |  | Experiment 1 | Experiment 2 |  |
|  |  | d 28 | d 11 | d 33 |
| without | 0 | 4.72 | 9.73<sup>a</sup> | 3.98 |
|  | 0.1 | 3.33 | 3.53<sup>b</sup> | 2.30 |
| with | 0 | 4.16 | 6.04<sup>ab</sup> | 4.61 |
|  | 0.1 | 3.38 | 4.13<sup>b</sup> | 2.80 |
| SEM<sup>1</sup> |  | 0.147 | 0.674 | 0.250 |
| Main effects |  |  |  |  |
| <u>Medication</u> |  |  |  |  |
| without |  | 4.02 | 6.63 | 3.14 |
| with |  | 3.77 | 5.08 | 3.70 |
| <u>β-glucanase (%)</u> |  |  |  |  |
| 0 |  | 4.44<sup>a</sup> | 7.89 | 4.29<sup>a</sup> |
| 0.1 |  | 3.35<sup>b</sup> | 3.83 | 2.55<sup>b</sup> |
| <u>Probability</u> |  |  |  |  |
| Medication |  | 0.25 | 0.11 | 0.17 |
| β-glucanase | | <.0001 | 0.0005 | 0.0002 |
| Medication × β-glucanase | | 0.16 | 0.03 | 0.86 |

<sup>a-b</sup>Means within a main effect or interaction not sharing a common superscript are significantly different ( $P \leq 0.05$ ).
<sup>1</sup>SEM - pooled standard error of mean (d 28; n=10 cages per treatment/ d 11; n=6 birds per treatment/ d 33; n=9 birds per treatment).

**Table 3.** Effects of diet medication and β-glucanase on ileal digesta short chain fatty acids of broiler chickens at 28 days of age (Experiment 1)

| Medication | BGase<sup>1</sup><br>(%) | SCFA μmol/g of wet ileal content | | | | | | | | | Molar percentage of total SCFA | | | | | | | |
| --- | --- | --- | --- | --- | --- | --- | --- | --- | --- | --- | --- | --- | --- | --- | --- | --- | --- | --- |
|  |  | Total | Ace | Pro | But | Isob | Val | Isov | Cap | Lac | Ace | Pro | But | Isob | Isov | Val | Cap | Lac |
| without | 0 | 165.8 | 61.8 | 22.2 | 10.6 | 2.7 | 3.3 | 2.9 | 1.3 | 60.6 | 37.5 | 13.1 | 6.4 | 1.6 | 1.7 | 1.9 | 0.7 | 36.6 |
|  | 0.1 | 157.2 | 59.1 | 20.8 | 10.3 | 2.9 | 2.6 | 2.2 | 1.0 | 58.0 | 37.6 | 13.3 | 6.5 | 1.8 | 1.4 | 1.6 | 0.6 | 36.9 |
| with | 0 | 173.5 | 66.4 | 23.4 | 10.8 | 2.5 | 2.7 | 2.9 | 1.5 | 63.0 | 38.3 | 13.2 | 6.3 | 1.4 | 1.6 | 1.5 | 0.8 | 36.5 |
|  | 0.1 | 156.9 | 59.1 | 21.8 | 10.3 | 2.4 | 2.6 | 2.6 | 1.2 | 56.5 | 37.6 | 14.0 | 6.6 | 1.4 | 1.6 | 1.6 | 0.8 | 36.1 |
| SEM<sup>2</sup> |  | 4.51 | 1.66 | 0.75 | 0.31 | 0.18 | 0.17 | 0.17 | 0.07 | 1.60 | 0.23 | 0.28 | 0.09 | 0.10 | 0.09 | 0.08 | 0.09 | 0.24 |
| Main effects |  |  |  |  |  |  |  |  |  |  |  |  |  |  |  |  |  |  |
| <u>Medication</u> |  |  |  |  |  |  |  |  |  |  |  |  |  |  |  |  |  |  |
| Without |  | 161.5 | 60.5 | 21.5 | 10.4 | 2.8 | 2.9 | 2.6 | 1.1 | 59.3 | 37.6 | 13.2 | 6.5 | 1.7 | 1.5 | 1.7 | 0.7 | 36.7 |
| With |  | 165.2 | 62.7 | 22.6 | 10.5 | 2.4 | 2.6 | 2.8 | 1.3 | 59.8 | 38.0 | 13.6 | 6.4 | 1.4 | 1.6 | 1.5 | 0.8 | 36.3 |
| <u>BGase (%)</u> |  |  |  |  |  |  |  |  |  |  |  |  |  |  |  |  |  |  |
| 0 |  | 169.6 | 64.1 | 22.8 | 10.7 | 2.6 | 3.0 | 2.9 | 1.4<sup>a</sup> | 61.8 | 37.9 | 13.2 | 6.3 | 1.5 | 1.7 | 1.7 | 0.8 | 36.6 |
| 0.1 |  | 157.0 | 59.1 | 21.3 | 10.3 | 2.6 | 2.6 | 2.4 | 1.1<sup>b</sup> | 57.2 | 37.6 | 13.6 | 6.6 | 1.6 | 1.5 | 1.6 | 0.7 | 36.5 |
| <u>Probability (%)</u> |  |  |  |  |  |  |  |  |  |  |  |  |  |  |  |  |  |  |
| Medication |  | 0.66 | 0.46 | 0.41 | 0.86 | 0.31 | 0.38 | 0.55 | 0.10 | 0.87 | 0.38 | 0.48 | 0.80 | 0.13 | 0.78 | 0.25 | 0.08 | 0.38 |
| BGase |  | 0.13 | 0.11 | 0.28 | 0.53 | 0.94 | 0.28 | 0.11 | 0.02 | 0.13 | 0.57 | 0.45 | 0.27 | 0.55 | 0.34 | 0.55 | 0.10 | 0.85 |
| Medication × BGase | | 0.63 | 0.45 | 0.94 | 0.90 | 0.67 | 0.36 | 0.52 | 0.73 | 0.51 | 0.40 | 0.59 | 0.61 | 0.57 | 0.34 | 0.23 | 0.35 | 0.47 |

<sup>a-d</sup>Means within a main effect or interaction not sharing a common superscript are significantly different ( $P \leq 0.05$ ).
<sup>1</sup>BGase - β-glucanase; SCFA - short chain fatty acids; Ace - Acetic acid; Pro - Propionic acid; But - Butyric acid; Isob - Isobutyric acid; Val - Valeric acid; Isov - Isovaleric acid; Cap - Caproic acid.
<sup>2</sup>SEM - pooled standard error of mean (n=20 birds per treatment).

**Table 4.**
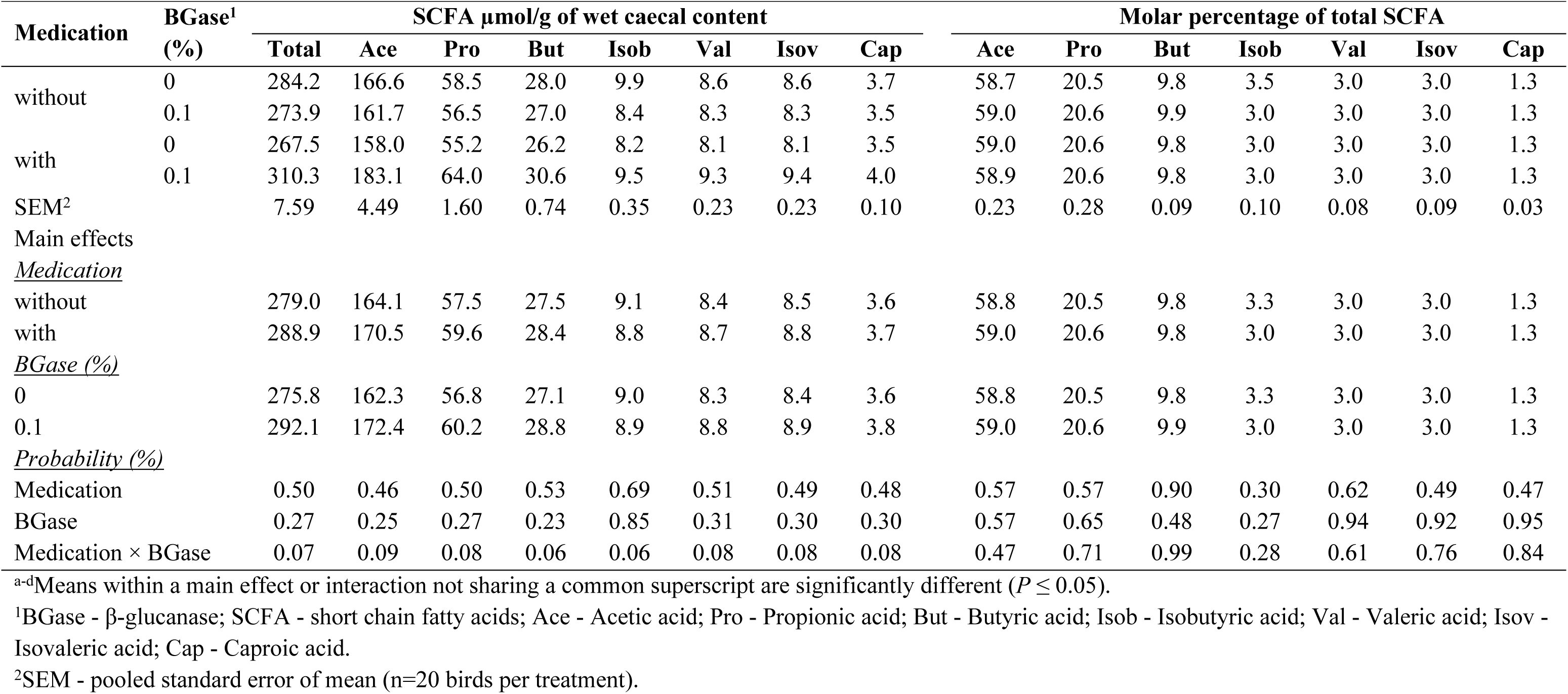
Effects of diet medication and β-glucanase on caecal short chain fatty acids of broiler chickens aged 28 days

To a large extent, dietary treatment did not affect ileal digesta SCFA of 11d old broilers in Experiment 2 (Table 6). The exception was a significant interaction between medication and BGase for valeric acid. Without medication, levels of valeric acid decreased with enzyme use, while levels increased with enzyme use when the medication was included in the diet. A similar trend (*P* = 0.10) was noted for isovaleric acid. Levels of caproic acid decreased with enzyme use. Interactions between the main effects were found for the molar percentages of valeric, isovaleric (*P* = 0.06), and caproic acids. In diets without medication, BGase did not affect acid concentration. When the medication was used, BGase increased acid levels. Dietary treatment interactions were also noted for the proportional levels of propionic and lactic acids. All mean differences were small and often not significant, but values tended to increase and decrease with BGase use in nonmedicated and medicated diets, respectively.

**Table 6.** Effects of diet medication and β-glucanase on ileal short chain fatty acids of broiler chickens aged 11 days (Experiment 2)

| Medication | BGase<sup>1</sup><br>(%) | SCFA μmol/g of wet ileal content |  |  |  |  |  |  |  | Molar percentage of total SCFA |  |  |  |  |  |  |
| --- | --- | --- | --- | --- | --- | --- | --- | --- | --- | --- | --- | --- | --- | --- | --- | --- |
|  |  | Total | Ace | Pro | But | Val | Isov | Cap | Lac | Ace | Pro | But | Val | Isov | Cap | Lac |
| without | 0 | 125.3 | 48.2 | 18.4 | 8.2 | 2.7<sup>a</sup> | 1.5 | 1.19 | 44.9 | 38.4 | 14.6<sup>ab</sup> | 6.5 | 2.1<sup>a</sup> | 1.2 | 0.9<sup>a</sup> | 35.8<sup>ab</sup> |
|  | 0.1 | 122.5 | 47.6 | 18.3 | 8.1 | 1.5<sup>bc</sup> | 1.4 | 0.79 | 44.6 | 38.8 | 14.9<sup>a</sup> | 6.6 | 1.2<sup>ab</sup> | 1.1 | 0.9<sup>a</sup> | 36.4<sup>ab</sup> |
| with | 0 | 121.5 | 46.8 | 18.0 | 7.6 | 1.3<sup>c</sup> | 1.4 | 1.19 | 45.1 | 38.6 | 14.8<sup>ab</sup> | 6.2 | 1.1<sup>b</sup> | 1.1 | 0.6<sup>b</sup> | 36.9<sup>a</sup> |
|  | 0.1 | 118.7 | 45.3 | 17.2 | 7.7 | 2.5<sup>ab</sup> | 2.5 | 1.10 | 42.1 | 38.2 | 14.5<sup>b</sup> | 6.5 | 2.1<sup>a</sup> | 2.1 | 0.9<sup>a</sup> | 35.4<sup>b</sup> |
| SEM<sup>2</sup> |  | 1.93 | 0.71 | 0.28 | 0.22 | 0.17 | 0.19 | 0.05 | 0.84 | 0.21 | 0.05 | 0.13 | 0.13 | 0.15 | 0.03 | 0.17 |
| Main effects |  |  |  |  |  |  |  |  |  |  |  |  |  |  |  |  |
| *Medication* |  |  |  |  |  |  |  |  |  |  |  |  |  |  |  |  |
| without |  | 123.9 | 47.9 | 18.3 | 8.2 | 2.1 | 1.4 | 0.99 | 44.8 | 38.6 | 14.8 | 6.6 | 1.7 | 1.1 | 0.7 | 36.1 |
| with |  | 120.6 | 46.1 | 17.6 | 7.6 | 1.9 | 1.9 | 1.14 | 43.6 | 38.4 | 14.6 | 6.4 | 1.6 | 1.6 | 0.9 | 36.2 |
| *BGase (%)* |  |  |  |  |  |  |  |  |  |  |  |  |  |  |  |  |
| 0 |  | 123.4 | 47.5 | 18.2 | 7.9 | 2.0 | 1.4 | 1.19<sup>a</sup> | 45.0 | 38.5 | 14.7 | 6.4 | 1.6 | 1.1 | 0.9 | 36.4 |
| 0.1 |  | 120.6 | 46.4 | 17.7 | 7.9 | 2.0 | 1.9 | 0.95<sup>b</sup> | 43.4 | 38.5 | 14.7 | 6.6 | 1.7 | 1.6 | 0.7 | 35.9 |
| *Probability (%)* |  |  |  |  |  |  |  |  |  |  |  |  |  |  |  |  |
| Medication |  | 0.29 | 0.16 | 0.16 | 0.24 | 0.53 | 0.17 | 0.10 | 0.45 | 0.64 | 0.22 | 0.42 | 0.69 | 0.12 | 0.02 | 0.89 |
| BGase |  | 0.43 | 0.41 | 0.39 | 0.99 | 0.98 | 0.17 | 0.01 | 0.30 | 0.94 | 0.79 | 0.54 | 0.77 | 0.13 | 0.01 | 0.16 |
| Medication × BGase |  | 0.99 | 0.75 | 0.50 | 0.90 | 0.0003 | 0.10 | 0.09 | 0.39 | 0.36 | 0.01 | 0.84 | 0.002 | 0.06 | 0.04 | 0.001 |

<sup>a-d</sup>Means within a main effect or interaction not sharing a common superscript are significantly different ( $P \leq 0.05$ ).
<sup>1</sup>BGase - β-glucanase; SCFA - short chain fatty acids; Ace - Acetic acid; Pro - Propionic acid; But - Butyric acid; Isob - Isobutyric acid; Val - Valeric acid; Isov - Isovaleric acid; Cap - Caproic acid.
<sup>2</sup>SEM - pooled standard error of mean (n=12 birds per treatment).

The interactions between medication and BGase use at 11 d were significant for total and individual caecal digesta SCFA (Table 7). The concentrations were higher with 0.1 compared to 0% BGase in the birds given diets without medication. However, BGase did not affect SCFA concentrations in the treatments with medication. Concentrations for birds fed medicated diets were lower than those fed un-medicated diets for the treatments with BGase. The molar percentages of propionic and isobutyric acids were decreased by medication, while enzyme use decreased and increased the proportions of acetic and butyric acids, and valeric acid, respectively. The interaction between main effects was significant for the proportional isovaleric levels, with enzyme tending to decrease levels in unmedicated diets and increase levels in medicated diets. Although the above effects were significant, differences were small.

**Table 7.** Effects of diet medication and β-glucanase on caecal short chain fatty acids of broiler chickens aged 11 days (Experiment 2)

| Medication | BGase<sup>1</sup><br>(%) | SCFA $\mu\text{mol/g}$ of wet caecal content | | | | | | | | Molar percentage of total SCFA | | | | | | |
| --- | --- | --- | --- | --- | --- | --- | --- | --- | --- | --- | --- | --- | --- | --- | --- | --- |
|  |  | Total | Ace | Pro | But | Isob | Val | Isov | Cap | Ace | Pro | But | Isob | Val | Isov | Cap |
| without | 0 | 228.6<sup>b</sup> | 134.1<sup>b</sup> | 49.7<sup>b</sup> | 22.7<sup>b</sup> | 7.4<sup>b</sup> | 4.3<sup>b</sup> | 7.4<sup>b</sup> | 2.7<sup>b</sup> | 58.6 | 21.8 | 9.9 | 3.2 | 1.7 | 3.3<sup>a</sup> | 0.1 |
|  | 0.1 | 306.6<sup>a</sup> | 176.5<sup>a</sup> | 66.3<sup>a</sup> | 30.0<sup>a</sup> | 9.9<sup>a</sup> | 9.7<sup>a</sup> | 9.8<sup>a</sup> | 4.2<sup>a</sup> | 57.5 | 21.6 | 9.7 | 3.2 | 3.1 | 3.2<sup>ab</sup> | 0.1 |
| with | 0 | 172.8<sup>b</sup> | 100.9<sup>bc</sup> | 36.4<sup>bc</sup> | 17.5<sup>b</sup> | 5.4<sup>c</sup> | 4.6<sup>b</sup> | 5.4<sup>c</sup> | 2.3<sup>b</sup> | 58.3 | 21.1 | 10.1 | 3.1 | 2.7 | 3.1<sup>b</sup> | 0.1 |
|  | 0.1 | 171.2<sup>b</sup> | 98.8<sup>c</sup> | 36.7<sup>c</sup> | 16.8<sup>b</sup> | 5.5<sup>c</sup> | 5.4<sup>b</sup> | 5.4<sup>c</sup> | 2.2<sup>b</sup> | 57.7 | 21.4 | 9.8 | 3.2 | 3.1 | 3.2<sup>ab</sup> | 0.1 |
| SEM<sup>2</sup> |  | 12.94 | 7.41 | 2.83 | 1.25 | 0.42 | 0.58 | 0.41 | 0.19 | 0.21 | 0.05 | 0.13 | 0.01 | 0.13 | 0.15 | 0.03 |
| Main effects |  |  |  |  |  |  |  |  |  |  |  |  |  |  |  |  |
| <u>Medication</u> |  |  |  |  |  |  |  |  |  |  |  |  |  |  |  |  |
| without |  | 267.6 | 155.3 | 58.0 | 26.3 | 8.7 | 7.0 | 8.6 | 3.4 | 58.1 | 21.7<sup>a</sup> | 9.8 | 3.2<sup>a</sup> | 2.4 | 3.2 | 0.1 |
| with |  | 172.0 | 99.8 | 36.6 | 17.2 | 5.4 | 5.0 | 5.4 | 2.3 | 58.0 | 21.3<sup>b</sup> | 9.8 | 3.1<sup>b</sup> | 2.9 | 3.1 | 0.1 |
| <u>BGase (%)</u> |  |  |  |  |  |  |  |  |  |  |  |  |  |  |  |  |
| 0 |  | 200.7 | 117.5 | 43.1 | 20.1 | 6.4 | 4.5 | 6.4 | 2.5 | 58.5<sup>a</sup> | 21.5 | 10.0<sup>a</sup> | 3.2 | 2.2<sup>b</sup> | 3.2 | 0.1 |
| 0.1 |  | 238.9 | 137.7 | 51.5 | 23.4 | 7.7 | 7.5 | 7.6 | 3.2 | 57.6<sup>b</sup> | 21.5 | 9.8<sup>b</sup> | 3.2 | 3.1<sup>a</sup> | 3.1 | 0.1 |
| <u>Probability (%)</u> |  |  |  |  |  |  |  |  |  |  |  |  |  |  |  |  |
| Medication |  | <.0001 | <.0001 | <.0001 | <.0001 | <.0001 | 0.02 | <.0001 | 0.0002 | 0.68 | 0.01 | 0.09 | 0.01 | 0.14 | 0.01 | 0.57 |
| BGase |  | 0.02 | 0.03 | 0.01 | 0.04 | 0.01 | 0.001 | 0.01 | 0.01 | 0.0004 | 0.91 | 0.01 | 0.89 | 0.007 | 0.93 | 0.34 |
| Medication × BGase | | 0.01 | 0.02 | 0.02 | 0.01 | 0.02 | 0.01 | 0.03 | 0.005 | 0.22 | 0.17 | 0.64 | 0.08 | 0.16 | 0.05 | 0.38 |

<sup>a-d</sup>Means within a main effect or interaction not sharing a common superscript are significantly different ( $P \leq 0.05$ ).
<sup>1</sup>BGase - β-glucanase; SCFA - short chain fatty acids; Ace - Acetic acid; Pro - Propionic acid; But - Butyric acid; Isob - Isobutyric acid; Val - Valeric acid; Isov - Isovaleric acid; Cap - Caproic acid.
<sup>2</sup>SEM - pooled standard error of mean (n=12 birds per treatment).

Medication and the interactions between medication and BGase did not affect the concentrations and molar percentages of ileal SCFA at d 33 (Table 8). All ileal SCFA concentrations except butyric acid were higher because of BGase use. In addition, the percentages of valeric and isovaleric acids were higher for the 0.1 compared to the 0% BGase treatment. In contrast, the lactic acid percentage was slightly lower with enzyme use.

**Table 8.** Effects of diet medication and β-glucanase on ileal short chain fatty acids of broiler chickens aged 33 days (Experiment 2)

| Medication | BGase<sup>1</sup><br>(%) | SCFA $\mu\text{mol/g}$ of wet ileal content | | | | | | | | Molar percentage of total SCFA | | | | | | |
| --- | --- | --- | --- | --- | --- | --- | --- | --- | --- | --- | --- | --- | --- | --- | --- | --- |
|  |  | Total | Ace | Pro | But | Val | Isov | Cap | Lac | Ace | Pro | But | Val | Isov | Cap | Lac |
| without | 0 | 115.2 | 44.6 | 17.0 | 7.6 | 1.5 | 1.6 | 1.0 | 41.6 | 38.7 | 14.79 | 6.6 | 1.3 | 1.4 | 0.8 | 36.1 |
|  | 0.1 | 125.0 | 47.8 | 18.1 | 8.1 | 2.6 | 2.7 | 1.1 | 44.3 | 38.2 | 14.52 | 6.5 | 2.1 | 2.1 | 0.9 | 35.4 |
| with | 0 | 118.9 | 46.0 | 17.5 | 7.8 | 1.7 | 1.9 | 1.0 | 42.7 | 38.7 | 14.74 | 6.6 | 1.4 | 1.6 | 0.8 | 35.9 |
|  | 0.1 | 123.0 | 47.1 | 17.9 | 7.5 | 2.6 | 2.6 | 1.1 | 43.8 | 38.3 | 14.60 | 6.1 | 2.1 | 2.1 | 0.9 | 35.6 |
| SEM<sup>2</sup> |  | 1.21 | 0.46 | 0.17 | 0.13 | 0.11 | 0.11 | 0.02 | 0.43 | 0.21 | 0.05 | 0.13 | 0.13 | 0.15 | 0.03 | 0.17 |
| Main effects |  |  |  |  |  |  |  |  |  |  |  |  |  |  |  |  |
| <u>Medication</u> |  |  |  |  |  |  |  |  |  |  |  |  |  |  |  |  |
| without |  | 120.1 | 46.2 | 17.6 | 7.8 | 2.1 | 2.1 | 1.0 | 42.9 | 38.5 | 14.6 | 6.5 | 1.7 | 1.7 | 0.9 | 35.7 |
| with |  | 120.9 | 46.5 | 17.7 | 7.7 | 2.2 | 2.3 | 1.0 | 43.2 | 38.5 | 14.6 | 6.3 | 1.8 | 1.8 | 0.9 | 35.7 |
| <u>BGase (%)</u> |  |  |  |  |  |  |  |  |  |  |  |  |  |  |  |  |
| 0 |  | 117.0<sup>b</sup> | 45.3<sup>b</sup> | 17.2<sup>b</sup> | 7.7 | 1.6<sup>b</sup> | 1.7<sup>b</sup> | 1.0<sup>b</sup> | 42.1<sup>b</sup> | 38.7 | 14.7 | 6.6 | 1.4<sup>b</sup> | 1.5<sup>b</sup> | 0.8 | 36.0<sup>a</sup> |
| 0.1 |  | 124.0<sup>a</sup> | 47.5<sup>a</sup> | 18.0<sup>a</sup> | 7.8 | 2.6<sup>a</sup> | 2.6<sup>a</sup> | 1.1<sup>a</sup> | 44.0<sup>a</sup> | 38.3 | 14.5 | 6.3 | 2.1<sup>a</sup> | 2.1<sup>a</sup> | 0.9 | 35.5<sup>b</sup> |
| <u>Probability (%)</u> |  |  |  |  |  |  |  |  |  |  |  |  |  |  |  |  |
| Medication |  | 0.73 | 0.72 | 0.69 | 0.51 | 0.71 | 0.48 | 0.88 | 0.70 | 0.91 | 0.88 | 0.30 | 0.77 | 0.53 | 0.82 | 0.91 |
| BGase |  | 0.003 | 0.02 | 0.02 | 0.68 | <.0001 | <.0001 | 0.01 | 0.02 | 0.30 | 0.10 | 0.12 | 0.001 | 0.003 | 0.18 | 0.001 |
| Medication × BGase | | 0.22 | 0.24 | 0.34 | 0.15 | 0.61 | 0.40 | 0.92 | 0.34 | 0.76 | 0.25 | 0.37 | 0.83 | 0.57 | 0.72 | 0.24 |

<sup>a-d</sup>Means within a main effect or interaction not sharing a common superscript are significantly different ( $P \leq 0.05$ ).
<sup>1</sup>BGase - β-glucanase, SCFA - short chain fatty acids; Ace - Acetic acid; Pro - Propionic acid; But - Butyric acid; Isob - Isobutyric acid; Val - Valeric acid; Isov - Isovaleric acid; Cap - Caproic acid.
<sup>2</sup>SEM - pooled standard error of mean (n=18 birds per treatment).

Main effect interactions were not found for the concentrations and molar percentages of caecal digesta SCFA at d 33 (Table 9). However, the concentrations of total SCFA and acetic acid were lower in medicated diets. Similarly, all other SCFA levels except butyric acid tended (*P* = 0.06-0.07) to be lower with medication use. The molar percentages of acetic acid decreased, while butyric, valeric (*P* = 0.08) and isovaleric (*P* = 0.09) acids increased with medication use. Enzyme use decreased the molar percentage of acetic acid and increased values for all other SCFA except butyric acid but minimal changes again, as noted earlier.

**Table 9.** Effects of diet medication and β-glucanase on caecal short chain fatty acids of broiler chickens aged 33 days (Experiment 2)

| Medication | BGase<sup>1</sup><br>(%) | SCFA $\mu\text{mol/g}$ of wet caecal content | | | | | | | | Molar percentage of total SCFA | | | | | | |
| --- | --- | --- | --- | --- | --- | --- | --- | --- | --- | --- | --- | --- | --- | --- | --- | --- |
|  |  | Total | Ace | Pro | But | Isob | Val | Isov | Cap | Ace | Pro | But | Isob | Val | Isov | Cap |
| without | 0 | 225.0 | 132.2 | 46.5 | 22.5 | 6.9 | 6.8 | 6.8 | 2.9 | 58.8 | 20.6 | 10.0 | 3.0 | 3.04 | 3.05 | 1.31 |
|  | 0.1 | 230.7 | 134.9 | 48.1 | 23.0 | 7.2 | 7.1 | 7.1 | 3.0 | 58.5 | 20.8 | 9.9 | 3.1 | 3.08 | 3.09 | 1.33 |
| with | 0 | 209.8 | 122.6 | 43.5 | 21.4 | 6.5 | 6.4 | 6.4 | 2.7 | 58.4 | 20.7 | 10.2 | 3.1 | 3.07 | 3.07 | 1.32 |
|  | 0.1 | 215.5 | 125.3 | 45.1 | 22.0 | 6.7 | 6.6 | 6.7 | 2.8 | 58.1 | 20.9 | 10.2 | 3.1 | 3.10 | 3.11 | 1.33 |
| SEM<sup>2</sup> |  | 3.78 | 2.17 | 0.82 | 0.38 | 0.12 | 0.12 | 0.12 | 0.05 | 0.21 | 0.05 | 0.13 | 0.01 | 0.13 | 0.15 | 0.03 |
| Main effects |  |  |  |  |  |  |  |  |  |  |  |  |  |  |  |  |
| <u>Medication</u> |  |  |  |  |  |  |  |  |  |  |  |  |  |  |  |  |
| without |  | 227.8<sup>a</sup> | 133.5<sup>a</sup> | 47.3 | 22.7 | 7.0 | 6.9 | 7.0 | 3.0 | 58.6<sup>a</sup> | 20.7 | 10.0<sup>b</sup> | 3.1 | 3.06 | 3.07 | 1.32 |
| with |  | 212.6<sup>b</sup> | 124.0<sup>b</sup> | 44.3 | 21.7 | 6.6 | 6.5 | 6.5 | 2.8 | 58.2<sup>b</sup> | 20.8 | 10.2<sup>a</sup> | 3.1 | 3.08 | 3.09 | 1.33 |
| <u>BGase (%)</u> |  |  |  |  |  |  |  |  |  |  |  |  |  |  |  |  |
| 0 |  | 217.4 | 127.4 | 45.0 | 22.0 | 6.7 | 6.6 | 6.6 | 2.8 | 58.6<sup>a</sup> | 20.7<sup>b</sup> | 10.1 | 3.0<sup>b</sup> | 3.05<sup>b</sup> | 3.06<sup>b</sup> | 1.31<sup>b</sup> |
| 0.1 |  | 223.1 | 130.1 | 46.6 | 22.5 | 6.9 | 6.8 | 6.9 | 2.9 | 58.3<sup>b</sup> | 20.9<sup>a</sup> | 10.0 | 3.1<sup>a</sup> | 3.09<sup>a</sup> | 3.09<sup>a</sup> | 1.33<sup>a</sup> |
| <u>Probability (%)</u> |  |  |  |  |  |  |  |  |  |  |  |  |  |  |  |  |
| Medication |  | 0.04 | 0.02 | 0.06 | 0.15 | 0.06 | 0.07 | 0.07 | 0.07 | 0.005 | 0.20 | 0.02 | 0.14 | 0.08 | 0.09 | 0.12 |
| BGase |  | 0.43 | 0.51 | 0.31 | 0.50 | 0.27 | 0.29 | 0.28 | 0.27 | 0.03 | 0.02 | 0.75 | 0.004 | 0.01 | 0.01 | 0.004 |
| Medication × BGase | | 0.99 | 0.99 | 0.99 | 0.93 | 0.98 | 0.99 | 0.99 | 0.94 | 0.93 | 0.97 | 0.85 | 0.88 | 0.93 | 0.97 | 0.59 |

<sup>a-d</sup>Means within a main effect or interaction not sharing a common superscript are significantly different ( $P \leq 0.05$ ).
<sup>1</sup>BGase - β-glucanase; SCFA - short chain fatty acids; Ace - Acetic acid; Pro - Propionic acid; But - Butyric acid; Isob - Isobutyric acid; Val - Valeric acid; Isov - Isovaleric acid; Cap - Caproic acid.
<sup>2</sup>SEM - pooled standard error of mean (n=18 birds per treatment).

Except for the duodenum, medication, BGase, and their interactions did not affect the digestive tract pH in Experiment 1 (Table 10). Enzyme use increased duodenal pH from 6.08 to 6.20. Main effect interactions were not found for the digestive tract pH, except for caecal pH at d 11 in Experiment 2 (Table 11); pH was lower with the enzyme use, but only in the diets without medication. Medication resulted in higher pH in the crop at d 11, and the ileum at both d 11 and 33. Duodenal and ileal pH was higher with the use of BGase at d 11. Gizzard and caecal pH were lower with the enzyme, and ileal pH was higher with the addition of diet BGase at d 33.

**Table 5.** Effects of diet medication and β-glucanase on gastro-intestinal pH of broiler chickens at day 28 (Experiment 1)

| Medication | β-glucanase (%) | Crop | Gizzard | Duodenum | Jejunum | Ileum | Caeca | Colon |
| --- | --- | --- | --- | --- | --- | --- | --- | --- |
| without | 0 | 5.29 | 3.54 | 6.05 | 5.99 | 7.08 | 6.02 | 6.92 |
|  | 0.1 | 5.23 | 3.26 | 6.19 | 6.01 | 7.26 | 6.04 | 7.17 |
| with | 0 | 5.43 | 3.23 | 6.10 | 5.96 | 7.25 | 5.90 | 7.08 |
|  | 0.1 | 5.20 | 3.17 | 6.21 | 6.05 | 7.27 | 5.93 | 7.13 |
| SEM<sup>1</sup> |  | 0.070 | 0.071 | 0.027 | 0.024 | 0.048 | 0.055 | 0.067 |
| Main effects |  |  |  |  |  |  |  |  |
| *Medication* |  |  |  |  |  |  |  |  |
| without |  | 5.26 | 3.40 | 6.12 | 5.99 | 7.17 | 6.03 | 7.04 |
| with |  | 5.31 | 3.20 | 6.16 | 6.00 | 7.26 | 5.91 | 7.11 |
| *β-glucanase (%)* |  |  |  |  |  |  |  |  |
| 0 |  | 5.36 | 3.39 | 6.08<sup>b</sup> | 5.97 | 7.16 | 5.96 | 7.00 |
| 0.1 |  | 5.21 | 3.22 | 6.20<sup>a</sup> | 6.03 | 7.26 | 5.98 | 7.15 |
| *Probability* |  |  |  |  |  |  |  |  |
| Medication |  | 0.70 | 0.15 | 0.46 | 0.89 | 0.25 | 0.29 | 0.61 |
| β-glucanase | | 0.29 | 0.21 | 0.01 | 0.16 | 0.20 | 0.82 | 0.22 |
| Medication × β-glucanase | | 0.55 | 0.41 | 0.80 | 0.40 | 0.29 | 0.94 | 0.43 |

<sup>a-b</sup>Means within a main effect or interaction not sharing a common superscript are significantly different ( $P \leq 0.05$ ).
<sup>1</sup>SEM - pooled standard error of mean (n=20 birds per treatment).

**Table 11.** Effects of diet medication and diet on gastro-intestinal pH of broiler chickens (Experiment 2)

| Medication | <sup>1</sup>BGase (%) | pH |  |  |  |  |  |  |  |  |  |  |  |
| --- | --- | --- | --- | --- | --- | --- | --- | --- | --- | --- | --- | --- | --- |
|  |  | d 11 |  |  |  |  |  | d 33 |  |  |  |  |  |
|  |  | Crop | Gizzard | Duodenum | Jejunum | Ileum | Caeca | Crop | Gizzard | Duodenum | Jejunum | Ileum | Caeca |
| without | 0 | 4.78 | 2.81 | 5.88 | 5.91 | 6.29 | 6.36<sup>a</sup> | 4.94 | 3.67 | 6.15 | 5.93 | 6.50 | 6.22 |
|  | 0.1 | 4.62 | 2.41 | 5.99 | 5.92 | 6.61 | 5.78<sup>b</sup> | 4.84 | 3.44 | 6.01 | 5.99 | 6.94 | 6.03 |
| with | 0 | 4.93 | 2.49 | 5.90 | 5.90 | 6.62 | 5.70<sup>b</sup> | 5.01 | 3.75 | 6.18 | 5.97 | 7.20 | 6.19 |
|  | 0.1 | 5.09 | 2.55 | 6.06 | 6.01 | 6.97 | 5.77<sup>b</sup> | 4.91 | 3.28 | 6.18 | 5.99 | 7.39 | 5.96 |
| SEM<sup>2</sup> |  | 0.052 | 0.057 | 0.024 | 0.018 | 0.053 | 0.061 | 0.052 | 0.057 | 0.024 | 0.018 | 0.053 | 0.061 |
| Main effects |  |  |  |  |  |  |  |  |  |  |  |  |  |
| *Medication* |  |  |  |  |  |  |  |  |  |  |  |  |  |
| without |  | 4.70<sup>b</sup> | 2.61 | 5.94 | 5.92 | 6.45<sup>b</sup> | 6.07 | 4.89 | 3.55 | 6.08 | 5.96 | 6.72<sup>b</sup> | 6.12 |
| with |  | 5.01<sup>a</sup> | 2.52 | 5.98 | 5.96 | 6.80<sup>a</sup> | 5.74 | 4.96 | 3.52 | 6.18 | 5.98 | 7.30<sup>a</sup> | 6.08 |
| *BGase (%)* |  |  |  |  |  |  |  |  |  |  |  |  |  |
| 0 |  | 4.85 | 2.65 | 5.89<sup>b</sup> | 5.91 | 6.45<sup>b</sup> | 6.03 | 4.97 | 3.71<sup>a</sup> | 6.16 | 5.95 | 6.85<sup>b</sup> | 6.21<sup>a</sup> |
| 0.1 |  | 4.86 | 2.48 | 6.03<sup>a</sup> | 5.97 | 6.79<sup>a</sup> | 5.78 | 4.87 | 3.36<sup>b</sup> | 6.09 | 5.99 | 7.17<sup>a</sup> | 5.99<sup>b</sup> |
| *Probability* |  |  |  |  |  |  |  |  |  |  |  |  |  |
| Medication |  | 0.001 | 0.41 | 0.33 | 0.25 | 0.0001 | 0.001 | 0.46 | 0.71 | 0.09 | 0.61 | <.0001 | 0.65 |
| BGase |  | 0.97 | 0.12 | 0.004 | 0.10 | 0.0002 | 0.01 | 0.29 | 0.001 | 0.22 | 0.28 | 0.0007 | 0.04 |
| Medication × BGase |  | 0.10 | 0.04 | 0.66 | 0.14 | 0.84 | 0.002 | 0.98 | 0.24 | 0.21 | 0.61 | 0.16 | 0.88 |

<sup>a-b</sup>Means within a main effect or interaction not sharing a common superscript are significantly different ( $P \leq 0.05$ ).
<sup>1</sup>BGase - β-glucanase; <sup>2</sup>SEM - pooled standard error of mean (d 11; n=12 birds per treatment, d 33; n=18 birds per treatment).

### Gastro-intestinal wall histomorphology

Gastrointestinal wall histology was examined only in Experiment 2 (Table 12). Treatment effects were not prevalent nor consistent between ages. At d 11, medication decreased the crypt depth, while β-glucanase decreased villi width. At 33 d, medication increased the number of acidic and decreased the number of mixed goblet cells per villus. The medication also increased the villi height to crypt depth ratio.

**Table 12.** Effects of medication and β-glucanase on histomorphology parameters in the ileum of broiler chickens (Experiment 2)

| Medication | BGase<sup>1</sup><br>(%) | d 11 |  |  |  |  |  |  | d 33 |  |  |  |  |  |  |
| --- | --- | --- | --- | --- | --- | --- | --- | --- | --- | --- | --- | --- | --- | --- | --- |
| | | Villi<br>height<br>(μm) | Villi<br>width<br>(μm) | Number of goblet<br>cells/villus | | | Crypt<br>depth<br>(μm) | Villi<br>height:<br>Crypt<br>depth | Villi<br>height<br>(μm) | Villi<br>width<br>(μm) | Number of goblet<br>cells/villus | | | Crypt<br>depth<br>(μm) | Villi<br>height:<br>Crypt<br>depth |
|  |  |  |  | Acidic | Neutral | Mixed |  |  |  |  | Acidic | Neutral | Mixed |  |  |
| without | 0 | 402 | 101 | 30 | 12 | 4 | 136 | 3.1 | 657 | 117 | 77 | 20 | 7 | 134 | 5 |
|  | 0.1 | 446 | 92 | 35 | 17 | 6 | 139 | 3.2 | 656 | 115 | 63 | 20 | 9 | 160 | 4 |
| with | 0 | 405 | 104 | 41 | 11 | 5 | 107 | 3.7 | 734 | 113 | 87 | 20 | 6 | 136 | 5 |
|  | 0.1 | 383 | 88 | 37 | 15 | 4 | 121 | 3.2 | 746 | 124 | 91 | 25 | 3 | 143 | 5 |
| SEM<sup>2</sup> |  | 22.27 | 2.20 | 2.59 | 1.30 | 0.46 | 5.21 | 0.19 | 23.26 | 2.60 | 4.44 | 1.74 | 0.96 | 4.61 | 0.18 |
| Main effects |  |  |  |  |  |  |  |  |  |  |  |  |  |  |  |
| <u>Medication</u> |  |  |  |  |  |  |  |  |  |  |  |  |  |  |  |
| without |  | 424 | 97 | 32 | 14 | 5 | 137<sup>a</sup> | 3.1 | 656 | 116 | 70<sup>b</sup> | 20 | 8<sup>a</sup> | 147 | 4<sup>b</sup> |
| with |  | 394 | 96 | 39 | 13 | 5 | 114<sup>b</sup> | 3.4 | 740 | 118 | 89<sup>a</sup> | 22 | 4<sup>b</sup> | 140 | 5<sup>a</sup> |
| <u>BGase (%)</u> |  |  |  |  |  |  |  |  |  |  |  |  |  |  |  |
| 0 |  | 404 | 102<sup>a</sup> | 35 | 11 | 5 | 121 | 3.4 | 695 | 115 | 82 | 20 | 6 | 135 | 5 |
| 0.1 |  | 414 | 90<sup>b</sup> | 36 | 16 | 5 | 130 | 3.2 | 701 | 120 | 77 | 22 | 6 | 151 | 4 |
| <u>Probability</u> |  |  |  |  |  |  |  |  |  |  |  |  |  |  |  |
| Medication |  | 0.54 | 0.91 | 0.21 | 0.56 | 0.82 | 0.01 | 0.41 | 0.07 | 0.62 | 0.03 | 0.48 | 0.04 | 0.39 | 0.03 |
| BGase |  | 0.83 | 0.01 | 0.96 | 0.08 | 0.96 | 0.32 | 0.58 | 0.90 | 0.29 | 0.52 | 0.51 | 0.98 | 0.06 | 0.13 |
| Medication × BGase | | 0.50 | 0.43 | 0.39 | 0.94 | 0.22 | 0.53 | 0.43 | 0.88 | 0.17 | 0.28 | 0.59 | 0.21 | 0.25 | 0.17 |

<sup>a-b</sup> Means within a main effect or interaction not sharing a common superscript are significantly different ( $P \leq 0.05$ ).
<sup>1</sup>BGase - β-glucanase.
<sup>2</sup>SEM - pooled standard error of mean (n=6 birds per treatment).

### Gastro-intestinal tract morphology

In Experiment 1, interactions were not found between BGase and medication for empty weights and lengths of the digestive tract sections, except for crop weight (Table 13). Crop weight was lower with enzyme use when the birds were fed a non-medicated diet, but the enzyme had no effect when the diets were medicated. Both ileum and colon weights were lower when the enzyme was fed. Crop content weight was higher, and duodenal and ileal content weights were lower when 0.1% BGase was fed (Table 14). Interactions were found for the content weights of the gizzard, jejunum and small intestine. The gizzard content weight tended to be higher and lower with enzyme use in birds fed non-medicated and medicated diets, respectively. Beta-glucanase resulted in lower jejunal and small intestinal content weights in the absence of dietary antibiotics but had no effect when the medication was used.

**Table 13.** Effects of diet medication and β-glucanase on gastro-intestinal tissue weights and lengths (proportional to body

| Medication | BGase<sup>1</sup><br>(%) | Empty weight (%) |  |  |  |  |  |  |  |  | Length (cm/100g) |  |  |  |  |  |
| --- | --- | --- | --- | --- | --- | --- | --- | --- | --- | --- | --- | --- | --- | --- | --- | --- |
|  |  | Crop | Proven | Gizzard | Duo | Jejunum | Ileum | SI | Caeca | Colon | Duo | Jejunum | Ileum | SI | Caeca | Colon |
| without | 0 | 0.34<sup>a</sup> | 0.38 | 1.20 | 0.73 | 1.37 | 1.00 | 3.08 | 0.36 | 0.17 | 1.73 | 4.22 | 4.18 | 10.07 | 1.67 | 0.41 |
|  | 0.1 | 0.29<sup>b</sup> | 0.38 | 1.32 | 0.73 | 1.30 | 0.91 | 2.94 | 0.37 | 0.14 | 1.75 | 4.01 | 4.11 | 9.87 | 1.69 | 0.39 |
| with | 0 | 0.30<sup>ab</sup> | 0.43 | 1.31 | 0.71 | 1.31 | 0.97 | 2.99 | 0.36 | 0.15 | 1.80 | 4.24 | 4.35 | 10.39 | 1.73 | 0.42 |
|  | 0.1 | 0.31<sup>ab</sup> | 0.38 | 1.33 | 0.74 | 1.28 | 0.93 | 2.94 | 0.37 | 0.15 | 1.79 | 4.23 | 4.29 | 10.28 | 1.68 | 0.42 |
| SEM<sup>2</sup> |  | 0.006 | 0.009 | 0.020 | 0.008 | 0.018 | 0.012 | 0.030 | 0.009 | 0.003 | 0.023 | 0.056 | 0.059 | 0.118 | 0.026 | 0.007 |
| Main effects |  |  |  |  |  |  |  |  |  |  |  |  |  |  |  |  |
| <u>Medication</u> |  |  |  |  |  |  |  |  |  |  |  |  |  |  |  |  |
| without |  | 0.32 | 0.38 | 1.26 | 0.73 | 1.33 | 0.96 | 3.01 | 0.36 | 0.16 | 1.74 | 4.12 | 4.15 | 9.97 | 1.68 | 0.40 |
| with |  | 0.30 | 0.40 | 1.32 | 0.73 | 1.30 | 0.95 | 2.97 | 0.37 | 0.15 | 1.79 | 4.23 | 4.32 | 10.33 | 1.71 | 0.42 |
| <u>BGase (%)</u> |  |  |  |  |  |  |  |  |  |  |  |  |  |  |  |  |
| 0 |  | 0.32 | 0.41 | 1.25 | 0.72 | 1.34 | 0.98<sup>a</sup> | 3.04 | 0.36 | 0.16<sup>a</sup> | 1.76 | 4.23 | 4.27 | 10.23 | 1.70 | 0.42 |
| 0.1 |  | 0.30 | 0.38 | 1.32 | 0.74 | 1.29 | 0.92<sup>b</sup> | 2.94 | 0.37 | 0.15<sup>b</sup> | 1.77 | 4.12 | 4.20 | 10.07 | 1.68 | 0.41 |
| <u>Probability</u> |  |  |  |  |  |  |  |  |  |  |  |  |  |  |  |  |
| Medication |  | 0.36 | 0.18 | 0.10 | 0.83 | 0.34 | 0.61 | 0.45 | 0.84 | 0.58 | 0.21 | 0.29 | 0.13 | 0.11 | 0.61 | 0.16 |
| BGase |  | 0.29 | 0.10 | 0.07 | 0.30 | 0.14 | 0.005 | 0.12 | 0.41 | 0.01 | 0.83 | 0.32 | 0.56 | 0.49 | 0.75 | 0.44 |
| Medication × BGase | | 0.007 | 0.13 | 0.18 | 0.47 | 0.57 | 0.31 | 0.40 | 0.98 | 0.08 | 0.75 | 0.35 | 0.97 | 0.82 | 0.48 | 0.64 |

<sup>a-b</sup>Means within a main effect or interaction not sharing a common superscript are significantly different ( $P \leq 0.05$ ).
<sup>1</sup>BGase - β-glucanase; Proven - proventriculus; Duo - duodenum; SI - small intestine.
<sup>2</sup>SEM - pooled standard error of mean (n=20 birds per treatment).

**Table 14.**
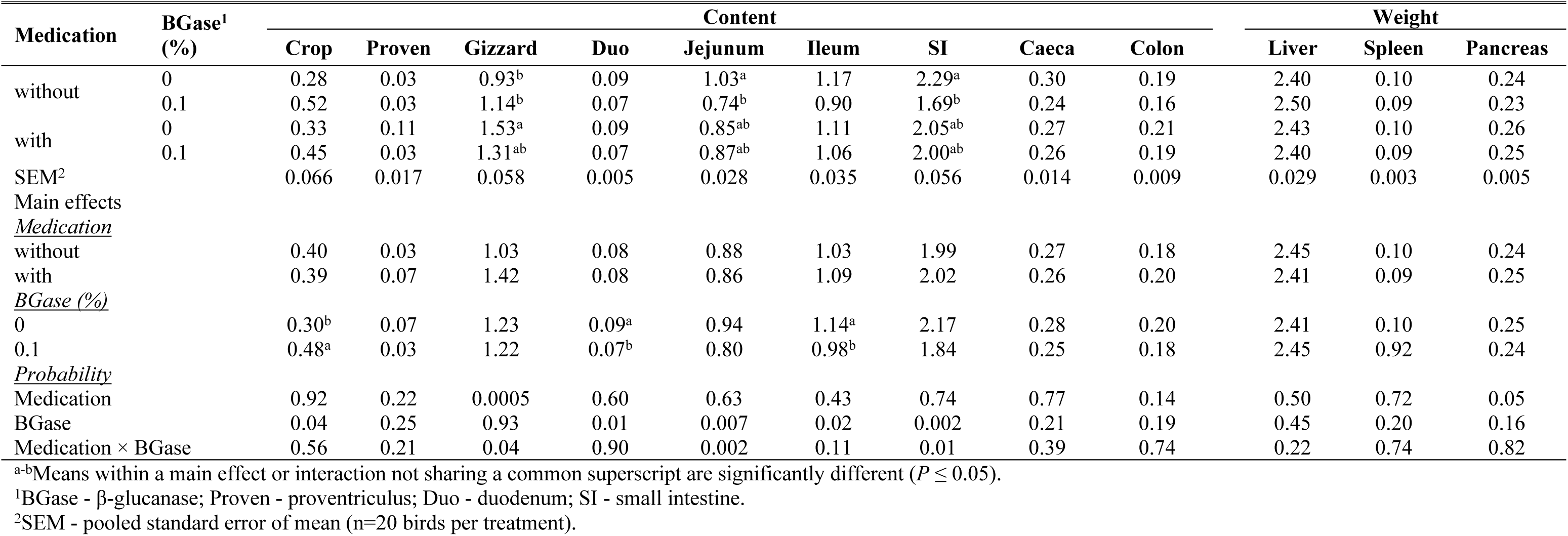
Effects of diet medication and β-glucanase on gastro-intestinal content and organ weights as a percentage of body weight of broiler chickens at d 28 (Experiment 1)

| Medication | BGase<sup>1</sup><br>(%) | Content |  |  |  |  |  |  |  |  | Weight |  |  |
| --- | --- | --- | --- | --- | --- | --- | --- | --- | --- | --- | --- | --- | --- |
|  |  | Crop | Proven | Gizzard | Duo | Jejunum | Ileum | SI | Caeca | Colon | Liver | Spleen | Pancreas |
| without | 0 | 0.28 | 0.03 | 0.93<sup>b</sup> | 0.09 | 1.03<sup>a</sup> | 1.17 | 2.29<sup>a</sup> | 0.30 | 0.19 | 2.40 | 0.10 | 0.24 |
|  | 0.1 | 0.52 | 0.03 | 1.14<sup>b</sup> | 0.07 | 0.74<sup>b</sup> | 0.90 | 1.69<sup>b</sup> | 0.24 | 0.16 | 2.50 | 0.09 | 0.23 |
| with | 0 | 0.33 | 0.11 | 1.53<sup>a</sup> | 0.09 | 0.85<sup>ab</sup> | 1.11 | 2.05<sup>ab</sup> | 0.27 | 0.21 | 2.43 | 0.10 | 0.26 |
|  | 0.1 | 0.45 | 0.03 | 1.31<sup>ab</sup> | 0.07 | 0.87<sup>ab</sup> | 1.06 | 2.00<sup>ab</sup> | 0.26 | 0.19 | 2.40 | 0.09 | 0.25 |
| SEM<sup>2</sup> |  | 0.066 | 0.017 | 0.058 | 0.005 | 0.028 | 0.035 | 0.056 | 0.014 | 0.009 | 0.029 | 0.003 | 0.005 |
| Main effects |  |  |  |  |  |  |  |  |  |  |  |  |  |
| <u>Medication</u> |  |  |  |  |  |  |  |  |  |  |  |  |  |
| without |  | 0.40 | 0.03 | 1.03 | 0.08 | 0.88 | 1.03 | 1.99 | 0.27 | 0.18 | 2.45 | 0.10 | 0.24 |
| with |  | 0.39 | 0.07 | 1.42 | 0.08 | 0.86 | 1.09 | 2.02 | 0.26 | 0.20 | 2.41 | 0.09 | 0.25 |
| <u>BGase (%)</u> |  |  |  |  |  |  |  |  |  |  |  |  |  |
| 0 |  | 0.30<sup>b</sup> | 0.07 | 1.23 | 0.09<sup>a</sup> | 0.94 | 1.14<sup>a</sup> | 2.17 | 0.28 | 0.20 | 2.41 | 0.10 | 0.25 |
| 0.1 |  | 0.48<sup>a</sup> | 0.03 | 1.22 | 0.07<sup>b</sup> | 0.80 | 0.98<sup>b</sup> | 1.84 | 0.25 | 0.18 | 2.45 | 0.92 | 0.24 |
| <u>Probability</u> |  |  |  |  |  |  |  |  |  |  |  |  |  |
| Medication |  | 0.92 | 0.22 | 0.0005 | 0.60 | 0.63 | 0.43 | 0.74 | 0.77 | 0.14 | 0.50 | 0.72 | 0.05 |
| BGase |  | 0.04 | 0.25 | 0.93 | 0.01 | 0.007 | 0.02 | 0.002 | 0.21 | 0.19 | 0.45 | 0.20 | 0.16 |
| Medication × BGase | | 0.56 | 0.21 | 0.04 | 0.90 | 0.002 | 0.11 | 0.01 | 0.39 | 0.74 | 0.22 | 0.74 | 0.82 |

<sup>a-b</sup>Means within a main effect or interaction not sharing a common superscript are significantly different ( $P \leq 0.05$ ).
<sup>1</sup>BGase - β-glucanase; Proven - proventriculus; Duo - duodenum; SI - small intestine.
<sup>2</sup>SEM - pooled standard error of mean (n=20 birds per treatment).

Interactions were found between medication and BGase for the empty proportional weights of the duodenum, jejunum, small intestine and caeca at d 11 (Table 15). For all segments, feeding diets without medication or enzyme resulted in the heaviest weights. Using an enzyme in nonmedicated diets reduced the segment weights (jejunum and small intestine), while enzyme use in diets with medication did not affect empty weight. Feeding an enzyme reduced the proventriculus empty weight. The length of the jejunum, ileum, small intestine and caeca were shorter with medication use. The dietary enzyme reduced the length of the jejunum and the small intestine. The content weight of the small intestine was lower, with the addition of BGase to the diets without medication (Table 16). Medication reduced the content weight of the crop and caeca, while BGase reduced the content weight of the gizzard, jejunum, ileum and colon. Diet medication reduced the pancreas weight, and diet enzyme increased liver weight and decreased pancreas weight.

**Table 15.** Effects of diet medication and β-glucanase on gastro-intestinal tissue weights and lengths (proportional to body weight) of broiler chickens at day 11

| Medication | BGase<sup>1</sup><br>(%) | Empty weight (%) |  |  |  |  |  |  |  |  | Length (cm/100g) |  |  |  |  |  |
| --- | --- | --- | --- | --- | --- | --- | --- | --- | --- | --- | --- | --- | --- | --- | --- | --- |
|  |  | Crop | Proven | Gizzard | Duodenum | Jejunum | Ileum | SI | Caeca | Colon | Duodenum | Jejunum | Ileum | SI | Caeca | Colon |
| without | 0 | 0.53 | 0.83 | 2.63 | 1.92<sup>a</sup> | 2.97<sup>a</sup> | 2.11 | 7.00<sup>a</sup> | 0.66<sup>a</sup> | 0.26 | 7.22 | 17.42 | 15.66 | 40.29 | 5.40 | 1.39 |
|  | 0.1 | 0.48 | 0.79 | 2.61 | 1.77<sup>ab</sup> | 2.63<sup>b</sup> | 1.88 | 6.27<sup>b</sup> | 0.60<sup>ab</sup> | 0.22 | 6.90 | 15.16 | 14.97 | 37.02 | 5.24 | 1.36 |
| with | 0 | 0.46 | 0.87 | 2.69 | 1.51<sup>b</sup> | 2.40<sup>b</sup> | 1.74 | 5.65<sup>c</sup> | 0.50<sup>b</sup> | 0.25 | 7.13 | 15.45 | 13.56 | 36.14 | 4.62 | 1.40 |
|  | 0.1 | 0.48 | 0.77 | 2.54 | 1.69<sup>ab</sup> | 2.67<sup>b</sup> | 1.78 | 6.13<sup>bc</sup> | 0.62<sup>ab</sup> | 0.25 | 6.11 | 14.64 | 13.75 | 34.49 | 4.99 | 1.34 |
| SEM<sup>2</sup> |  | 0.018 | 0.018 | 0.043 | 0.039 | 0.053 | 0.043 | 0.109 | 0.020 | 0.006 | 0.219 | 0.273 | 0.329 | 0.584 | 0.121 | 0.035 |
| Main effects |  |  |  |  |  |  |  |  |  |  |  |  |  |  |  |  |
| <u>Medication</u> |  |  |  |  |  |  |  |  |  |  |  |  |  |  |  |  |
| without |  | 0.50 | 0.81 | 2.62 | 1.84 | 2.80 | 2.00<sup>a</sup> | 6.64 | 0.63 | 0.24 | 7.06 | 16.29<sup>a</sup> | 15.31<sup>a</sup> | 38.65<sup>a</sup> | 5.32<sup>a</sup> | 1.37 |
| with |  | 0.47 | 0.82 | 2.62 | 1.60 | 2.54 | 1.76<sup>b</sup> | 5.89 | 0.56 | 0.25 | 6.62 | 15.05<sup>b</sup> | 13.65<sup>b</sup> | 35.31<sup>b</sup> | 4.80<sup>b</sup> | 1.37 |
| <u>BGase (%)</u> |  |  |  |  |  |  |  |  |  |  |  |  |  |  |  |  |
| 0 |  | 0.49 | 0.85<sup>a</sup> | 2.66 | 1.72 | 2.69 | 1.93 | 6.33 | 0.58 | 0.25 | 7.17 | 16.43<sup>a</sup> | 14.61 | 38.21<sup>a</sup> | 5.01 | 1.39 |
| 0.1 |  | 0.48 | 0.78<sup>b</sup> | 2.58 | 1.73 | 2.65 | 1.83 | 6.20 | 0.61 | 0.24 | 6.50 | 14.90<sup>b</sup> | 14.36 | 35.76<sup>b</sup> | 5.12 | 1.35 |
| <u>Probability</u> |  |  |  |  |  |  |  |  |  |  |  |  |  |  |  |  |
| Medication |  | 0.16 | 0.77 | 0.92 | 0.0009 | 0.001 | 0.003 | <.0001 | 0.07 | 0.44 | 0.26 | 0.004 | 0.01 | 0.001 | 0.03 | 0.92 |
| BGase |  | 0.70 | 0.04 | 0.29 | 0.90 | 0.62 | 0.19 | 0.42 | 0.43 | 0.11 | 0.09 | 0.0007 | 0.69 | 0.01 | 0.65 | 0.41 |
| Medication × BGase |  | 0.15 | 0.42 | 0.41 | 0.02 | 0.0004 | 0.08 | 0.0005 | 0.01 | 0.15 | 0.36 | 0.08 | 0.48 | 0.40 | 0.26 | 0.74 |

<sup>a-b</sup>Means within a main effect or interaction not sharing a common superscript are significantly different ( $P \leq 0.05$ ).
<sup>1</sup>BGase - β-glucanase; Proven - proventriculus; SI - small intestine.
<sup>2</sup>SEM - pooled standard error of mean (n=12 birds per treatment).

**Table 16.** Effects of diet medication and β-glucanase on gastro-intestinal content and organ weights as a percentage of body weight of broiler chickens at day 11

| Medication | BGase<sup>1</sup><br>(%) | Content |  |  |  |  |  |  |  |  | Weight |  |  |
| --- | --- | --- | --- | --- | --- | --- | --- | --- | --- | --- | --- | --- | --- |
|  |  | Crop | Proventriculus | Gizzard | Duodenum | Jejunum | Ileum | SI | Caeca | Colon | Liver | Spleen | Pancreas |
| without | 0 | 0.48 | 0.06 | 0.89 | 0.08 | 0.59 | 0.60 | 1.26<sup>a</sup> | 0.08 | 0.06 | 4.05 | 0.13 | 0.57 |
|  | 0.1 | 0.54 | 0.05 | 0.81 | 0.05 | 0.45 | 0.41 | 0.89<sup>c</sup> | 0.11 | 0.04 | 4.74 | 0.11 | 0.50 |
| with | 0 | 0.29 | 0.11 | 0.99 | 0.05 | 0.53 | 0.51 | 1.08<sup>b</sup> | 0.07 | 0.07 | 4.19 | 0.13 | 0.50 |
|  | 0.1 | 0.37 | 0.06 | 0.73 | 0.04 | 0.45 | 0.44 | 0.93<sup>bc</sup> | 0.07 | 0.05 | 4.48 | 0.12 | 0.49 |
| SEM<sup>2</sup> |  | 0.035 | 0.008 | 0.034 | 0.006 | 0.018 | 0.018 | 0.727 | 0.006 | 0.004 | 0.070 | 0.004 | 0.011 |
| Main effects |  |  |  |  |  |  |  |  |  |  |  |  |  |
| *Medication* |  |  |  |  |  |  |  |  |  |  |  |  |  |
| without |  | 0.51<sup>a</sup> | 0.05 | 0.85 | 0.06 | 0.52 | 0.50 | 1.08 | 0.09<sup>a</sup> | 0.05 | 4.39 | 0.12 | 0.53<sup>a</sup> |
| with |  | 0.33<sup>b</sup> | 0.08 | 0.86 | 0.04 | 0.49 | 0.47 | 1.00 | 0.07<sup>b</sup> | 0.06 | 4.34 | 0.12 | 0.50<sup>b</sup> |
| *BGase (%)* |  |  |  |  |  |  |  |  |  |  |  |  |  |
| 0 |  | 0.38 | 0.08 | 0.94<sup>a</sup> | 0.06 | 0.56<sup>a</sup> | 0.55<sup>a</sup> | 1.17 | 0.07 | 0.07<sup>a</sup> | 4.12<sup>b</sup> | 0.13 | 0.54<sup>a</sup> |
| 0.1 |  | 0.46 | 0.05 | 0.77<sup>b</sup> | 0.04 | 0.45<sup>b</sup> | 0.42<sup>b</sup> | 0.91 | 0.09 | 0.04<sup>b</sup> | 4.61<sup>a</sup> | 0.11 | 0.50<sup>b</sup> |
| *Probability* |  |  |  |  |  |  |  |  |  |  |  |  |  |
| Medication |  | 0.008 | 0.08 | 0.89 | 0.09 | 0.29 | 0.36 | 0.11 | 0.03 | 0.09 | 0.63 | 0.64 | 0.04 |
| BGase |  | 0.26 | 0.08 | 0.009 | 0.06 | <.0001 | 0.0001 | <.0001 | 0.20 | 0.005 | 0.0002 | 0.10 | 0.03 |
| Medication × BGase |  | 0.85 | 0.15 | 0.15 | 0.22 | 0.16 | 0.06 | 0.02 | 0.22 | 0.91 | 0.09 | 0.57 | 0.13 |

<sup>a-b</sup>Means within a main effect or interaction not sharing a common superscript are significantly different ( $P \leq 0.05$ ).
<sup>1</sup>BGase - β-glucanase; SI - small intestine
<sup>2</sup>SEM - pooled standard error of mean (n=12 birds per treatment).

Diet medication decreased the empty proportional weights of the duodenum, jejunum, ileum, small intestine and colon, and decreased the lengths of the same digestive tract segments in 33 d old broilers (Table 17). Dietary BGase resulted in lower empty weights for the crop, ileum and small intestine; enzyme also reduced the lengths of the duodenum and ileum. Interactions between the main effects were found for the empty jejunum weight, and the lengths of the jejunum and small intestine. For the interactions, enzyme use resulted in smaller tissues when non-medicated diets were fed, but had no effect when diets contained medication. Medication resulted in smaller digestive tract segments in these interactions.

**Table 17.** Effects of diet medication and β-glucanase on gastro-intestinal tissue weights and lengths (proportional to body weight) of broiler chickens at day 33

| Medication | BGase<sup>1</sup><br>(%) | Empty weight (%) |  |  |  |  |  |  |  |  | Length (cm/100g) |  |  |  |  |  |
| --- | --- | --- | --- | --- | --- | --- | --- | --- | --- | --- | --- | --- | --- | --- | --- | --- |
|  |  | Crop | Proven | Gizzard | Duo | Jejunum | Ileum | SI | Caeca | Colon | Duo | Jejunum | Ileum | SI | Caeca | Colon |
| without | 0 | 0.30 | 0.38 | 1.12 | 0.87 | 1.64<sup>a</sup> | 1.13 | 3.64 | 0.37 | 0.17 | 1.80 | 4.49<sup>a</sup> | 4.42 | 10.70<sup>a</sup> | 0.63 | 0.41 |
|  | 0.1 | 0.29 | 0.39 | 1.23 | 0.86 | 1.53<sup>a</sup> | 1.00 | 3.38 | 0.38 | 0.15 | 1.63 | 3.88<sup>b</sup> | 3.86 | 9.37<sup>b</sup> | 0.71 | 0.38 |
| with | 0 | 0.33 | 0.44 | 1.14 | 0.71 | 1.24<sup>b</sup> | 0.98 | 2.92 | 0.35 | 0.15 | 1.57 | 3.43<sup>c</sup> | 3.36 | 8.35<sup>c</sup> | 0.60 | 0.32 |
|  | 0.1 | 0.27 | 0.36 | 1.16 | 0.70 | 1.28<sup>b</sup> | 0.92 | 2.90 | 0.37 | 0.15 | 1.47 | 3.40<sup>c</sup> | 3.34 | 8.20<sup>c</sup> | 0.64 | 0.35 |
| SEM<sup>2</sup> |  | 0.006 | 0.015 | 0.022 | 0.014 | 0.029 | 0.018 | 0.051 | 0.008 | 0.004 | 0.029 | 0.078 | 0.089 | 0.172 | 0.031 | 0.010 |
| Main effects |  |  |  |  |  |  |  |  |  |  |  |  |  |  |  |  |
| <u>Medication</u> |  |  |  |  |  |  |  |  |  |  |  |  |  |  |  |  |
| without |  | 0.29 | 0.38 | 1.17 | 0.86<sup>a</sup> | 1.58 | 1.06<sup>a</sup> | 3.51<sup>a</sup> | 0.38 | 0.16<sup>a</sup> | 1.71<sup>a</sup> | 4.19 | 4.14<sup>a</sup> | 10.03 | 0.67 | 0.40<sup>a</sup> |
| with |  | 0.30 | 0.40 | 1.15 | 0.70<sup>b</sup> | 1.26 | 0.95<sup>b</sup> | 2.91<sup>b</sup> | 0.36 | 0.15<sup>b</sup> | 1.52<sup>b</sup> | 3.41 | 3.35<sup>b</sup> | 8.27 | 0.62 | 0.33<sup>b</sup> |
| <u>BGase (%)</u> |  |  |  |  |  |  |  |  |  |  |  |  |  |  |  |  |
| 0 |  | 0.31<sup>a</sup> | 0.41 | 1.13 | 0.79 | 1.44 | 1.05<sup>a</sup> | 3.28<sup>a</sup> | 0.36 | 0.16 | 1.68<sup>a</sup> | 3.96 | 3.89<sup>a</sup> | 9.52 | 0.62 | 0.37 |
| 0.1 |  | 0.28<sup>b</sup> | 0.38 | 1.20 | 0.78 | 1.40 | 0.96<sup>b</sup> | 3.14<sup>b</sup> | 0.38 | 0.15 | 1.55<sup>b</sup> | 3.64 | 3.60<sup>b</sup> | 8.78 | 0.67 | 0.36 |
| <u>Probability</u> |  |  |  |  |  |  |  |  |  |  |  |  |  |  |  |  |
| Medication |  | 0.80 | 0.57 | 0.62 | <.0001 | <.0001 | 0.0005 | <.0001 | 0.36 | 0.01 | 0.0003 | <.0001 | <.0001 | <.0001 | 0.15 | 0.0004 |
| BGase |  | 0.005 | 0.27 | 0.12 | 0.55 | 0.33 | 0.005 | 0.04 | 0.22 | 0.11 | 0.01 | 0.009 | 0.04 | 0.004 | 0.11 | 0.88 |
| Medication × BGase | | 0.12 | 0.20 | 0.31 | 0.83 | 0.05 | 0.28 | 0.10 | 0.88 | 0.15 | 0.47 | 0.01 | 0.06 | 0.02 | 0.68 | 0.09 |

<sup>a-c</sup>Means within a main effect or interaction not sharing a common superscript are significantly different ( $P \leq 0.05$ ).
<sup>1</sup>BGase - β-glucanase; Proven - proventriculus; Duo - duodenum; SI - small intestine.
<sup>2</sup>SEM - pooled standard error of mean (n=18 birds per treatment).

The content weights of the duodenum and colon decreased with the use of BGase at d 33 (Table 18). Medication similarly decreased the content weight of the duodenum. Interactions between medication and enzyme were found for the content weights of the gizzard (*P* = 0.06), jejunum, ileum, small intestine and colon (*P* = 0.06). For the jejunum, ileum, small intestine and colon segments, enzyme reduced weights in non-medicated diets but did not affect content weights in the presence of medication. For gizzard content weights, enzyme increased and decreased values in diets without and with medication, respectively. An interaction was also found for liver weight. The largest weight was found for the birds fed diets with no medication or enzyme; the addition of enzyme to the unmedicated diet resulted in lower weight, and the liver weights for medicated diets were smallest and unaffected by enzyme in the diet.

**Table 18.** Effects of diet medication and β-glucanase on gastro-intestinal content and organ weights as a percentage of body weight of broiler chickens at day 33

| Medication | BGase<sup>1</sup><br>(%) | Content |  |  |  |  |  |  |  |  | Weight |  |  |
| --- | --- | --- | --- | --- | --- | --- | --- | --- | --- | --- | --- | --- | --- |
|  |  | Crop | Proventriculus | Gizzard | Duodenum | Jejunum | Ileum | SI | Caeca | Colon | Liver | Spleen | Pancreas |
| without | 0 | 1.54 | 0.11 | 1.18 | 0.12 | 1.31<sup>a</sup> | 1.49<sup>a</sup> | 2.91<sup>a</sup> | 0.27 | 0.23 | 3.16<sup>a</sup> | 0.12 | 0.27 |
|  | 0.1 | 1.44 | 0.06 | 1.33 | 0.09 | 0.86<sup>b</sup> | 0.97<sup>b</sup> | 1.91<sup>b</sup> | 0.32 | 0.14 | 2.88<sup>b</sup> | 0.12 | 0.27 |
| with | 0 | 1.46 | 0.34 | 1.56 | 0.08 | 1.03<sup>b</sup> | 1.12<sup>b</sup> | 2.21<sup>b</sup> | 0.25 | 0.17 | 2.57<sup>c</sup> | 0.12 | 0.26 |
|  | 0.1 | 1.11 | 0.07 | 1.24 | 0.07 | 0.95<sup>b</sup> | 0.91<sup>b</sup> | 1.92<sup>b</sup> | 0.27 | 0.17 | 2.58<sup>c</sup> | 0.12 | 0.26 |
| SEM<sup>2</sup> |  | 0.096 | 0.043 | 0.060 | 0.006 | 0.039 | 0.050 | 0.084 | 0.015 | 0.011 | 0.040 | 0.004 | 0.005 |
| Main effects |  |  |  |  |  |  |  |  |  |  |  |  |  |
| <u>Medication</u> |  |  |  |  |  |  |  |  |  |  |  |  |  |
| without |  | 1.49 | 0.09 | 1.26 | 0.10<sup>a</sup> | 1.08 | 1.23 | 2.41 | 0.29 | 0.18 | 3.02 | 0.12 | 0.27 |
| with |  | 1.28 | 0.20 | 1.40 | 0.07<sup>b</sup> | 0.99 | 1.02 | 2.07 | 0.26 | 0.17 | 2.57 | 0.12 | 0.26 |
| <u>BGase (%)</u> |  |  |  |  |  |  |  |  |  |  |  |  |  |
| 0 |  | 1.50 | 0.23 | 1.37 | 0.10<sup>a</sup> | 1.17 | 1.31 | 2.56 | 0.26 | 0.20<sup>a</sup> | 2.86 | 0.12 | 0.26 |
| 0.1 |  | 1.27 | 0.06 | 1.29 | 0.08<sup>b</sup> | 0.90 | 0.94 | 1.91 | 0.29 | 0.15<sup>b</sup> | 2.73 | 0.12 | 0.26 |
| <u>Probability</u> |  |  |  |  |  |  |  |  |  |  |  |  |  |
| Medication |  | 0.28 | 0.16 | 0.22 | 0.006 | 0.15 | 0.009 | 0.01 | 0.21 | 0.61 | <.0001 | 0.54 | 0.13 |
| BGase |  | 0.24 | 0.06 | 0.46 | 0.02 | 0.0002 | <.0001 | <.0001 | 0.20 | 0.03 | 0.01 | 0.93 | 0.81 |
| Medication × BGase | | 0.52 | 0.19 | 0.06 | 0.37 | 0.006 | 0.04 | 0.007 | 0.52 | 0.06 | 0.01 | 0.93 | 0.90 |

<sup>a-c</sup>Means within a main effect or interaction not sharing a common superscript are significantly different ( $P \leq 0.05$ ).
<sup>1</sup>BGase - β-glucanase; SI - small intestine
<sup>2</sup>SEM - pooled standard error of mean (n=18 birds per treatment).

### Performance parameters

Interactions between medication and BGase were significant or nearly significant for BWG and FI from 0-7 d, 7-14 d (*P* = 0.06) and 0-28 d (*P* = 0.06-0.07), and F:G from 0-7 d in Experiment 1 (Table 19). Body weight gain and FI followed a similar response to treatments. In birds fed diets without medication, the addition of BGase tended to reduce gain or feed consumption, however in those fed diets with medication, enzyme either did not affect or increased these response criteria. For the 0-7 d F:G ratio interaction, enzyme decreased and increased feed efficiency in unmedicated diets and medicated diets, respectively.

**Table 19.** Effects of diet medication and β-glucanase on performance parameters of broiler chickens (Experiment 1)

| Medication | β-glucanase (%) | BWG<sup>1</sup> (kg) | | | | | FI (kg) | | | | | F:G | | | | |
| --- | --- | --- | --- | --- | --- | --- | --- | --- | --- | --- | --- | --- | --- | --- | --- | --- |
|  |  | d | d | d | d | d | d | d | d | d | d | d | d | d | d | d |
|  |  | 0-7 | 7-14 | 14-21 | 21-28 | 0-28 | 0-7 | 7-14 | 14-21 | 21-28 | 0-28 | 0-7 | 7-14 | 14-21 | 21-28 | 0-28 |
| without | 0 | 0.143<sup>a</sup> | 0.303 | 0.507 | 0.699 | 1.650 | 0.167<sup>a</sup> | 0.421 | 0.729 | 1.055 | 2.371 | 1.17<sup>b</sup> | 1.39 | 1.44 | 1.53 | 1.45 |
|  | 0.1 | 0.126<sup>c</sup> | 0.296 | 0.498 | 0.656 | 1.575 | 0.157<sup>b</sup> | 0.399 | 0.705 | 1.004 | 2.265 | 1.26<sup>a</sup> | 1.35 | 1.42 | 1.54 | 1.44 |
| with | 0 | 0.130<sup>bc</sup> | 0.284 | 0.492 | 0.668 | 1.573 | 0.160<sup>ab</sup> | 0.387 | 0.706 | 1.000 | 2.251 | 1.23<sup>a</sup> | 1.36 | 1.44 | 1.50 | 1.43 |
|  | 0.1 | 0.135<sup>ab</sup> | 0.301 | 0.494 | 0.677 | 1.607 | 0.160<sup>ab</sup> | 0.409 | 0.695 | 1.012 | 2.275 | 1.19<sup>b</sup> | 1.36 | 1.41 | 1.50 | 1.42 |
| SEM<sup>2</sup> |  | 1.562 | 2.966 | 4.564 | 10.050 | 14.222 | 1.172 | 4.887 | 5.856 | 11.406 | 18.375 | 0.008 | 0.011 | 0.009 | 0.014 | 0.007 |
| Main effects |  |  |  |  |  |  |  |  |  |  |  |  |  |  |  |  |
| <u>Medication</u> |  |  |  |  |  |  |  |  |  |  |  |  |  |  |  |  |
| Without |  | 0.134 | 0.299 | 0.503 | 0.678 | 1.612 | 0.162 | 0.410 | 0.717 | 1.030 | 2.318 | 1.21 | 1.37 | 1.43 | 1.53 | 1.45 |
| With |  | 0.132 | 0.292 | 0.493 | 0.673 | 1.591 | 0.160 | 0.398 | 0.700 | 1.006 | 2.263 | 1.21 | 1.36 | 1.42 | 1.50 | 1.43 |
| <u>β-glucanase (%)</u> |  |  |  |  |  |  |  |  |  |  |  |  |  |  |  |  |
| 0 |  | 0.136 | 0.293 | 0.500 | 0.684 | 1.612 | 0.163 | 0.404 | 0.717 | 1.027 | 2.311 | 1.20 | 1.38 | 1.44 | 1.52 | 1.44 |
| 0.1 |  | 0.130 | 0.298 | 0.496 | 0.666 | 1.591 | 0.159 | 0.404 | 0.700 | 1.008 | 2.270 | 1.22 | 1.35 | 1.41 | 1.52 | 1.43 |
| <u>Probability</u> |  |  |  |  |  |  |  |  |  |  |  |  |  |  |  |  |
| Medication |  | 0.36 | 0.21 | 0.32 | 0.79 | 0.43 | 0.35 | 0.17 | 0.15 | 0.29 | 0.12 | 0.70 | 0.55 | 0.69 | 0.21 | 0.12 |
| β-glucanase | | 0.01 | 0.38 | 0.71 | 0.36 | 0.45 | 0.04 | 0.99 | 0.14 | 0.39 | 0.25 | 0.06 | 0.30 | 0.20 | 0.96 | 0.26 |
| Medication × β-glucanase | | <.0001 | 0.06 | 0.54 | 0.17 | 0.06 | 0.02 | 0.06 | 0.55 | 0.17 | 0.07 | <.0001 | 0.44 | 0.85 | 0.90 | 0.85 |

<sup>a-c</sup>Means within a main effect or interaction not sharing a common superscript are significantly different ( $P \leq 0.05$ ).
<sup>1</sup>BWG - body weight gain; FI - feed intake; F:G - feed to gain ratio.
<sup>2</sup>SEM - pooled standard error of mean (n=10 cages per treatment).

In Experiment 2, interactions between main effects were significant for BWG for all periods (Table 20), but the nature of the response changed with age. From 0-11 d, medication increased gain, while enzyme did not affect gain in birds fed diets without medication and tended to increase gain in the medicated diet. Weight gain from 11 to 22 d was increased by enzyme regardless of diet medication. From 22-32 d, enzyme increased gain in the non-medicated diets but had no effect when diets contain medication. Overall, weight gain (0-32 d) was increased by enzyme use, regardless of diet medication but to a greater extent in the absence of medication.

**Table 20.** Effects of diet medication and β-glucanase on performance parameters of broiler chickens vaccinated for coccidiosis (Experiment 2)

| Medication | BGase<sup>1</sup><br>(%) | BWG (kg) |  |  |  | FI (kg) |  |  |  | F:G |  |  |  |
| --- | --- | --- | --- | --- | --- | --- | --- | --- | --- | --- | --- | --- | --- |
|  |  | d | d | d | d | d | d | d | d | d | d | d | d |
|  |  | 0-11 | 11-22 | 22-32 | 0-32 | 0-11 | 11-22 | 22-32 | 0-32 | 0-11 | 11-22 | 22-32 | 0-32 |
| without | 0 | 0.243<sup>b</sup> | 0.562<sup>d</sup> | 0.788<sup>c</sup> | 1.594<sup>d</sup> | 0.328 | 0.979 | 1.540<sup>bc</sup> | 2.846 | 1.321<sup>b</sup> | 1.617<sup>a</sup> | 1.939<sup>a</sup> | 1.721<sup>a</sup> |
|  | 0.1 | 0.236<sup>b</sup> | 0.622<sup>c</sup> | 0.881<sup>b</sup> | 1.740<sup>c</sup> | 0.331 | 0.982 | 1.499<sup>c</sup> | 2.813 | 1.372<sup>a</sup> | 1.471<sup>b</sup> | 1.688<sup>b</sup> | 1.561<sup>b</sup> |
| with | 0 | 0.262<sup>a</sup> | 0.675<sup>b</sup> | 0.963<sup>a</sup> | 1.900<sup>b</sup> | 0.331 | 1.049 | 1.581<sup>ab</sup> | 2.961 | 1.242<sup>c</sup> | 1.429<sup>b</sup> | 1.627<sup>c</sup> | 1.497<sup>c</sup> |
|  | 0.1 | 0.270<sup>a</sup> | 0.702<sup>a</sup> | 0.981<sup>a</sup> | 1.954<sup>a</sup> | 0.339 | 1.071 | 1.588<sup>a</sup> | 2.998 | 1.236<sup>c</sup> | 1.423<sup>b</sup> | 1.593<sup>c</sup> | 1.479<sup>c</sup> |
| SEM<sup>2</sup> |  | 0.002 | 0.640 | 0.904 | 0.025 | 0.002 | 0.008 | 0.009 | 0.017 | 0.011 | 0.015 | 0.024 | 0.017 |
| Main effects |  |  |  |  |  |  |  |  |  |  |  |  |  |
| *Medication* |  |  |  |  |  |  |  |  |  |  |  |  |  |
| without |  | 0.240 | 0.591 | 0.835 | 1.667 | 0.329<sup>b</sup> | 0.981<sup>b</sup> | 1.520 | 2.829<sup>b</sup> | 1.347 | 1.544 | 1.813 | 1.641 |
| with |  | 0.266 | 0.689 | 0.972 | 1.927 | 0.335<sup>a</sup> | 1.060<sup>a</sup> | 1.584 | 2.905<sup>a</sup> | 1.239 | 1.426 | 1.610 | 1.488 |
| *BGase (%)* |  |  |  |  |  |  |  |  |  |  |  |  |  |
| 0 |  | 0.252 | 0.618 | 0.876 | 1.747 | 0.329<sup>b</sup> | 1.014 | 1.560 | 2.904 | 1.282 | 1.523 | 1.783 | 1.609 |
| 0.1 |  | 0.253 | 0.662 | 0.931 | 1.847 | 0.335<sup>a</sup> | 1.027 | 1.544 | 2.905 | 1.304 | 1.447 | 1.641 | 1.520 |
| *Probability* |  |  |  |  |  |  |  |  |  |  |  |  |  |
| Medication |  | <.0001 | <.0001 | <.0001 | <.0001 | 0.01 | <.0001 | <.0001 | <.0001 | <.0001 | <.0001 | <.0001 | <.0001 |
| BGase |  | 0.77 | <.0001 | <.0001 | <.0001 | 0.01 | 0.18 | 0.14 | 0.92 | 0.01 | <.0001 | <.0001 | <.0001 |
| Medication × BGase | | 0.006 | 0.02 | 0.001 | 0.002 | 0.29 | 0.33 | 0.04 | 0.06 | 0.001 | <.0001 | <.0001 | <.0001 |

<sup>a-c</sup>Means within a main effect or interaction not sharing a common superscript are significantly different ( $P \leq 0.05$ ).
<sup>1</sup>BGase - β-glucanase; BWG - body weight gain; FI - feed intake; F:G - feed to gain ratio.
<sup>2</sup>SEM - pooled standard error of mean (n=9 pens per treatment).

Medication and enzyme use increased FI from 0-11 d, and medication similarly increased FI from 11-22 d. Interactions between medication and enzyme were significant from 22-32 d and approached significance (*P* = 0.06) for the overall experiment. In both cases, the use of dietary BGase tended to decrease FI when medication was not fed and increase FI when it was.

Interactions were found between medication and BGase for F:G in all periods. Medication increased feed efficiency throughout the trial, but as was the case for BWG, the nature of the interaction with enzyme use changed with bird age. During the 0-11 d period, F:G increased with enzyme use when birds were fed non-medicated diets, but had no effect when the medication was used. For the remainder of the periods, including the total trial, enzyme decreased F:G in birds fed non-medicated diets, but had no effect in broilers consuming medicated diets.

The total mortality of the trials was 3.8 and 3.9% in Experiment 1 and 2, respectively, and not affected by HB or BGase. In Experiment 2, the mortality attributed to coccidiosis (by necropsy) was identified as 4.3% of the total mortality. However, 46.7% of the total mortality was detected as systemic infection, including necrotic enteritis. Subclinical coccidiosis in the birds may damage the intestinal epithelial membrane and thereby enhance systemic infections due to bacterial translocation. It can be concluded that the vaccination with Coccivac-B52 induced a disease challenge in the birds from Experiment 2 according to the detailed analysis of mortality data.

## Discussion

With minor exceptions, all three molecular weight parameters for soluble ileal digesta β-glucan were lower with the enzyme use, which confirms exogenous BGase mediates the depolymerization of HB β-glucan in broiler chickens. In addition, the reduction of MW-10% with BGase in both experiments further supports β-glucan depolymerization since it demonstrates the increased proportion of small molecular weight soluble β-glucan in ileal digesta. Overall, the response for Mp was similar in both experiments, which indicates that β-glucan depolymerization is independent of the disease status and the age of the animal. Further, Mw from Experiment 1 also supports the reduction of molecular weight in the ileal digesta soluble β-glucan with the use of BGase. In contrast, BGase increased Mw at both ages in Experiment 2 in the treatments with antibiotics. The reason for the increased β-glucan Mw is unknown but could relate to the aggregation of smaller weight β-glucan molecules [42–44] or enzyme mediated release of higher molecular weight, insoluble β-glucan that had not yet been depolymerized. The reduction of β-glucan molecular weight and the increased proportion of small molecular weight soluble β-glucan encourage the assessment of performance and digestive tract characteristics through increased β-glucan fermentation in broilers. Further, the proportion of small molecular weight β-glucan might be a critical assessment in chickens since chicken microbiota preferred small molecular sugars and peptides over complex polysaccharides and proteins in a study that investigated the utilization of nutrients by chicken caecal and human faecal microbes using an *in vitro* assay [45].

The molecular weight values were numerically lower at d 33 compared to d 11 in Experiment 2, which might be associated with an age-related adaptation of gut microbiota to utilize fibre [46]. Further, molecular weight parameters were lower in Experiment 1 compared to both ages in Experiment 2. Although the experiments cannot be compared statistically, it does draw attention to experimental variation. The analyses of samples were completed at three different times. However, the probability that analytical error accounted for the variation is unlikely because the determination of β-glucan molecular weight distribution using size exclusion chromatography and Calcofluor post-column derivatization is a well-established technique in food science [38] and all laboratory work was completed in the same lab by the senior author. A more plausible explanation for the difference relates to variation in β-glucan characteristics in the barley samples that were fed. The birds were fed diets containing CDC Fibar in both experiments; however, the samples were different in the two experiments. Although they were the same cultivar, environmental conditions such as germination may have impacted β-glucan molecular weight. High moisture content in the environment might activate endogenous enzymes in barley and degrade non-starch polysaccharides, including β-glucan, which is supported by the improved nutritive value of barley with water treatment [47]. Moreover, the molecular weight differences in the two experiments could be attributed to the resident gut microbiota being markedly different between the studies that could harbor different β-glucanase capabilities. The variable gut microbiota composition among the broiler chickens derived from the same breeder flock and raised under the same conditions, including diets, support the difference related to the microbial enzyme activity [48]. To the best of our knowledge, this, and other research in the same lab, are the first to document molecular weight changes in soluble β-glucan because of exogenous BGase use in chickens fed barley diets. The BGase effect on the reduction of ileal soluble β-glucan molecular weight in this study is in agreement with previous results from our lab [49].

The molecular weight parameters in the two experiments decreased with medication when there was no added BGase in the diet, which is an unexpected finding since the medication does not contain endo-β-glucanase activity. It is possibly due to the effect of the antibiotics on modification of the gastro-intestinal microbial population [50–52] resulting in microbiota with an increased capacity to degrade high molecular weight β-glucan into low molecular weight polysaccharides and oligosaccharides. *In vitro* studies have demonstrated that strict anaerobic caecal microbiota, including *Bacteroides ovatus*, *B. uniformis*, *B. capillosus, Enterococcus faecium*, *Clostridium perfringens* and *Streptococcus* strains in broiler chickens are capable of degrading mixed-linked β-glucan [53]. The changes in the intestinal microbial populations were also reported in pigs fed barley-based diets [54, 55], although they were not the same bacterial species as in the above-mentioned *in vitro* assays. However, medication was not able to breakdown high molecular weight β-glucan to the same extent as BGase. It demonstrates the higher efficacy of feed BGase in comparison to the BGase originated from microbiota in the chicken gastro-intestinal tract of degrading high molecular weight β-glucan.

Exogenous BGase depolymerizes high molecular weight soluble β-glucan into low molecular weight β-glucan in the ileal digesta, which leads to a reduction of ileal viscosity in broiler chickens, and this is evident in both experiments. However, the medication did not affect ileal viscosity in broiler chickens, although the molecular weight was reduced with the addition of antibiotics to the broiler diets. Nevertheless, viscosity changes at 11 d are similar to the molecular weight data where enzyme decreased values, but the decrease was smaller with medicated diets, and it is primarily due to the low viscosity in the treatments with medication when BGase was not used. In addition to the degree of polymerization of β-glucan, there might be other factors that affect intestinal viscosity, including the concentration, solubility, structure, and configuration of non-starch polysaccharides including β-glucan and arabinoxylan in the digesta [56–58]. Further, medication might have shifted the ileal microbial population in a way that leads to increased intestinal mucus production, which can contribute to ileal viscosity [59, 60]. In addition, high amounts of NSP in the diet also increase intestinal mucus production in monogastric animals [59, 61].

Overall, BGase reduced the empty weights, lengths, and content weights in the digestive tract segments in both experiments, which agrees with previous broiler research that used the same diets but without medication [62]. The reduction in size coincides with increased digestive efficiency associated with enzyme use and has been reported previously [63, 64]. In addition, the reduction of gastro-intestinal content weights might be associated with increased feed passage rate in the gastro-intestinal tract [65, 66] since exogenous BGase decreases digesta viscosity and thereby increases digestive function in the broiler chickens [67, 68]. Further, HB mediated larger digestive tract might hold more digesta that leads to increased gastro-intestinal content weights in the current study. Medication decreased the empty weights and lengths from the duodenum to colon and the content weights of the digestive tract segments. The reduction of digestive tract size and content follows previous research that observed the decreased intestinal tract weights and lengths with in-feed antibiotics (Bacitracin methylene disalicylate and virginiamycin) in broiler chickens [69]. The use of specific antibiotics in feed reduces the growth of pathogenic bacteria in the digestive tract of chickens through the modification of microbial diversity and relative abundance, and immune status [29, 30], and thereby increases nutrient digestibility. The reduction of relative abundance of gut microbiota reduces the competition with the host and thus enables the host to extract all the required nutrients, and thereby the digestive tract size might be reduced [70, 71]. Further, diet medication might increase nutrient digestion due to increased utilization of non-starch polysaccharides by the gut microbiota by selecting for a more effective fibre degrading microbiome, which is supported by β-glucan molecular weight reduction with antibiotics addition to the diets in the current research. The effects of medication on relative digestive tract size and content weights were mostly significant when the HB based diets did not contain BGase since the enzyme also decreases digestive tract size by increasing nutrient digestibility in broiler chickens.

Levels of SCFA and pH in the digestive tract were used to estimate the effects of diet BGase and antibiotics on carbohydrate fermentation. Diet BGase and medication depolymerized soluble β-glucan in HB in the ileal digesta of broiler chickens, which may influence carbohydrate fermentation in the lower digestive tract. Ileal pH was higher with BGase use at both ages of broiler chickens in Experiment 2. A BGase mediated increase in ileal pH is contradictory to the current hypothesis of an enzyme-dependent enhancement of carbohydrate fermentation that might be expected based on a large quantity of low molecular weight β-glucan resulting from high molecular weight β-glucan depolymerization due to enzyme use. The increased ileal pH might relate to the increased feed passage rate from the ileum to caeca with the reduction of soluble β-glucan molecular weight, which permits less time for the bacterial fermentation in the ileum [49]. However, ileal pH is contradictory to total, and individual SCFA concentrations in the ileum since BGase increased SCFA levels at d 33 in the current study. A reduction of caecal pH with enzyme (d 11 without medication; d 33) might indicate increased carbohydrate fermentation in the caeca, which is in agreement with previous research [25]. Further, BGase increased SCFA concentrations in the caeca (d 11 without medication) in the current study, which corresponds with the caecal pH at d 11. Overall, the results suggest BGase has shifted bacterial fermentation from the ileum to caeca in broiler chickens.

The antibiotic-induced modification of the gastro-intestinal microbial population might affect the production of SCFA, which in turn influences the enzyme response on carbohydrate fermentation in broiler chickens. Medication affected intestinal pH in a similar fashion to BGase, and similar to the findings of [28], who found increased ileal pH and lowered caecal pH with the addition of salinomycin and Zn bacitracin to broiler diets. However, diet medication did not affect the concentrations of SCFA in the ileum, whereas it decreased total and most of the individual SCFA concentrations in the caeca in the current study, which is again contradictory to the caecal pH. The reduction of caecal pH might be due to the effect of antibiotics on reducing protein putrefaction to a greater extent than it did SCFA production in the caeca. However, the concentrations of alkalizing metabolites, including the biogenic amines, are not available in the current study. Nevertheless, the reduction of caecal SCFA concentration was in accordance to the study completed by [72] that used salinomycin in broiler feed. Antibiotics modulate the microbial population of the chicken digestive tract [73, 74], and these microbes might not effectively utilize the fermentable fibre, including β-glucan in the chicken digestive tract due to the lower production of microbial-derived non-starch polysaccharidases. However, it is contradictory to the findings of the ileal β-glucan molecular weight distribution, since medication reduced the molecular weight, which demonstrates the presence of gastro-intestinal bacteria that could secrete non-starch polysaccharidases. The resulting SCFA might have been immediately utilized by gut microbes to produce other metabolic products and affects the measured levels of SCFA. Of note, the crop pH was higher with diet medication. The crop is colonized by BGase-secreting microbiota [75], and medication modifies the crop microbiota, thereby affects carbohydrate fermentation [76].

A few treatment main effects were found for ileal histo-morphology, but no interactions were significant. Medication increased villus height to crypt depth ratio in the ileum, which is an indication of increased nutrient absorption surface [77] that eventually leads to the enhancement of nutrient digestion and performance of chickens. The effect of diet medication on reducing digestive tract size and content also supports the increased nutrient digestibility, which is indicated by the higher villus height to crypt depth ratio. In addition, medication decreased crypt depth in the ileum. Increased crypt depth indicates high cell proliferation in the intestinal epithelial cells [78], which is an indication of inflammation in the intestinal mucosa. Thus the mucosa enhances healing from the inflammatory damage by increasing cell proliferation [79, 80]. Inflammation is a protective mechanism, although uncontrolled and chronic inflammation may damage the affected tissues [81, 82]. Therefore, the reduction of crypt depth is considered as a positive entity that enhances bird health. The use of specific diet medication shifts bacterial distribution in the digestive tract of broiler chickens towards saccharolytic fermentation [31] and increases SCFA production, including butyrate, that could increase digestive tract epithelial growth [83], and it might be the cause for the high villus height: crypt depth in the ileum. Short chain fatty acids, especially butyrate, have the potential to affect inflammation by regulating inflammatory cytokines [84, 85]. However, the medication did not affect total SCFA or butyrate in the ileum in the current research.

Treatment affected SCFA concentrations, and intestinal pH in coccidiosis challenged broiler chickens, but not in battery-cage raised and unchallenged birds. Further, the treatment effects were larger for broilers at 11 d (mostly infected with *Eimeria* spp) compared to the same birds at 33 d (mostly recovered from the disease) in the coccidiosis challenge study. *Eimeria* spp disturbs the lower gastro-intestinal microbial population in broilers [86, 87] due to the epithelial damage of the intestinal mucosa, and this, in turn, affects SCFA production [88]. On the other hand, a precise estimate of SCFA production might not be measured in the current study due to the limitations of the digesta collection procedure. Partial absorption of SCFA to the portal circulation before sample collection, which leads to under-estimation of the values, and ileal and caecal evacuation that is affected by the time of the sample collection, results in individual bird variability in results. In addition, protein fermentation affects digesta pH since some of the protein fermentation products, including ammonia, indoles, phenols and biogenic amines, increase pH in the digestive tract of chickens [89].

Performance variables were within the normal range, according to Ross 308 Broiler Performance Objectives [34]. The interaction between BGase and medication was significant for BWG and F:G at all the periods of the broiler production cycle in Experiment 2. Over the entire experiment, medication increased both BWG and feed efficiency of broilers. However, the medication response was higher without the use of BGase since exogenous BGase positively influence growth performance in the current study. Both Zn Bacitracin and ionophore anticoccidials have been classified as growth-promoting drugs in broiler chickens due to their positive impact on body weight gain and feed efficiency [28,90,91]. Antibiotics in the diets shift the gastro-intestinal microbial population towards a diversified and potentially beneficial microbiota [29, 92]. Among the beneficial changes is an increase in carbohydrate fermentation [93], including β-glucan, and positively affect gastro-intestinal physiology and health and helps in improving the production performance of broiler chickens. Short chain fatty acids, especially butyric acid, produced as a result of carbohydrate fermentation, increase energy supply to intestinal epithelial cells [94], increases nutrient absorptive surface area by increasing villi size [95, 96], and also decreases harmful pathogenic bacteria in the lower digestive tract of chickens [93]. Villi height to crypt depth ratio in the ileum increased with medication in the current study, which supports antibiotics mediated enhancement of the ileal absorptive surface area in broiler chickens. However, total and individual SCFA concentrations in the caeca decreased with the addition of antibiotics, which is contradictory to carbohydrate fermentation induced improvement of physiological and growth parameters in the current research.

Beta-glucanase increased the BWG and feed efficiency of broiler chickens after d 11, although these parameters were lower with the use of BGase before d 11. These results agree with previous research that used same diets without medication, which observed the poor production performance in young birds (< 11 d) [97]. The poor performance of younger birds may be attributed to an undesirable effect of the increased quantity of low molecular weight carbohydrates on the gut microbiota due to the coccidiosis challenge-induced diseased state and the immature status of the digestive system and gut microbiota. In the study of [97], BGase dosage of 0.01% increased broiler performance for the same age period (0 to 11 days) when compared to 0% BGase. However, 0.1% BGase did not affect the BWG and reduced the feed efficiency in the birds aged < 11 d but increased these parameters after d 11. Moreover, BGase decreased the total requirement of medication in HB-based diets in terms of achieving a high production performance, as the medication response on performance variables decreased with the addition of BGase to the diets. It demonstrates the ability of BGase to partially replace diet medication in HB-based diets to feed broiler chickens. In contrast to the results of Experiment 2, the effects of medication and BGase on performance variables were not significant in the production cycle except the period of d 0-7 of broiler chickens in Experiment 1, where birds were grown in battery cages without disease challenge. The environment of battery cages is relatively hygienic compared to litter floor pens and is generally considered to present less pathogenic bacterial exposure with the birds. It might be the reason for less significant effects of medication and enzyme on production parameters in the battery cage study.

In conclusion, feed BGase and medication can depolymerize high molecular weight soluble β-glucan of HB into low molecular weight β-glucan in the digestive tract of broilers in both experiments; however, the response was higher with BGase compared to medication. The effects of diet medication and BGase on carbohydrate fermentation were not consistent across sample collections in the two experiments according to SCFA levels and intestinal pH, although treatment effects were observed in certain instances. Exogenous BGase and medication increased the growth performance of broiler chickens. Moreover, BGase reduced the necessity of antibiotics and anticoccidials in HB-based diets to achieve a high level of production performance of broiler chickens challenged for coccidiosis.

## Acknowledgments

The authors would like to acknowledge the Poultry Centre staff at the University of Saskatchewan, and Dawn Abbott and Tracy Exley for their technical support.

## References

1. Castanon JIR. History of the use of antibiotic as growth promoters in European poultry feeds. Poult Sci. 2007;86: 2466–2471. doi:10.3382/ps.2007-00249

2. Gadde U, Kim WH, Oh ST, Lillehoj HS. Alternatives to antibiotics for maximizing growth performance and feed efficiency in poultry: a review. Anim Health Res Rev. 2017;18: 26– 45. doi:10.1017/S1466252316000207

3. Nhung NT, Chansiripornchai N, Carrique-Mas JJ. Antimicrobial resistance in bacterial poultry pathogens: a review. Front Vet Sci. 2017;4. doi:10.3389/fvets.2017.00126

4. Lillehoj H, Liu Y, Calsamiglia S, Fernandez-Miyakawa ME, Chi F, Cravens RL, et al. Phytochemicals as antibiotic alternatives to promote growth and enhance host health. Vet Res. 2018;49: 76. doi:10.1186/s13567-018-0562-6

5. Suresh G, Das RK, Brar SK, Rouissi T, Ramirez AA, Chorfi Y, et al. Alternatives to antibiotics in poultry feed: molecular perspectives. Crit Rev Microbiol. 2018;44: 318–335. doi:10.1080/1040841X.2017.1373062

6. Allen HK, Levine UY, Looft T, Bandrick M, Casey TA. Treatment, promotion, commotion: antibiotic alternatives in food-producing animals. Trends Microbiol. 2013;21: 114–119. doi:10.1016/j.tim.2012.11.001

7. Seal BS, Lillehoj HS, Donovan DM, Gay CG. Alternatives to antibiotics: a symposium on the challenges and solutions for animal production. Anim Health Res Rev. 2013;14: 78–87. doi:10.1017/S1466252313000030

8. Gaggìa F, Mattarelli P, Biavati B. Probiotics and prebiotics in animal feeding for safe food production. Int J Food Microbiol. 2010;141 Suppl 1: S15–28. doi:10.1016/j.ijfoodmicro.2010.02.031

9. Baurhoo B, Phillip L, Ruiz-Feria CA. Effects of purified lignin and mannan oligosaccharides on intestinal integrity and microbial populations in the ceca and litter of broiler chickens. Poult Sci. 2007;86: 1070–1078. doi:10.1093/ps/86.6.1070

10. Baurhoo B, Ferket PR, Zhao X. Effects of diets containing different concentrations of mannanoligosaccharide or antibiotics on growth performance, intestinal development, cecal and litter microbial populations, and carcass parameters of broilers. Poult Sci. 2009;88: 2262–2272. doi:10.3382/ps.2008-00562

11. Xu ZR, Hu CH, Xia MS, Zhan XA, Wang MQ. Effects of dietary fructooligosaccharide on digestive enzyme activities, intestinal microflora and morphology of male broilers. Poult Sci. 2003;82: 1030–1036. doi:10.1093/ps/82.6.1030

12. Shang Y, Regassa A, Kim JH, Kim WK. The effect of dietary fructooligosaccharide supplementation on growth performance, intestinal morphology, and immune responses in broiler chickens challenged with *Salmonella* Enteritidis lipopolysaccharides. Poult Sci. 2015;94: 2887–2897. doi:10.3382/ps/pev275

13. Flickinger EA, Van Loo J, Fahey GC. Nutritional responses to the presence of inulin and oligofructose in the diets of domesticated animals: a review. Crit Rev Food Sci Nutr. 2003;43: 19–60. doi:10.1080/10408690390826446

14. Bogusławska-Tryk M, Piotrowska A, Burlikowska K. Dietary fructans and their potential beneficial influence on health and performance parameters in broiler chickens. J Cent Eur Agric. 2012;13: 272–291. doi:https://doi.org/10.5513/JCEA01/13.2.1045

15. Kim G-B, Seo YM, Kim CH, Paik IK. Effect of dietary prebiotic supplementation on the performance, intestinal microflora, and immune response of broilers. Poult Sci. 2011;90: 75–82. doi:10.3382/ps.2010-00732

16. Khodambashi Emami N, Samie A, Rahmani HR, Ruiz-Feria CA. The effect of peppermint essential oil and fructooligosaccharides, as alternatives to virginiamycin, on growth performance, digestibility, gut morphology and immune response of male broilers. Anim Feed Sci Technol. 2012;175: 57–64. doi:10.1016/j.anifeedsci.2012.04.001

17. Courtin CM, Swennen K, Broekaert WF, Swennen Q, Buyse J, Decuypere E, et al. Effects of dietary inclusion of xylooligosaccharides, arabinoxylooligosaccharides and soluble arabinoxylan on the microbial composition of caecal contents of chickens. J Sci Food Agric. 2008;88: 2517–2522. doi:10.1002/jsfa.3373

18. De Maesschalck C, Eeckhaut V, Maertens L, De Lange L, Marchal L, Nezer C, et al. Effects of xylo-oligosaccharides on broiler chicken performance and microbiota. Appl Environ Microbiol. 2015;81: 5880–5888. doi:10.1128/AEM.01616-15

19. Eeckhaut V, Van Immerseel F, Dewulf J, Pasmans F, Haesebrouck F, Ducatelle R, et al. Arabinoxylooligosaccharides from wheat bran inhibit *Salmonella* colonization in broiler chickens. Poult Sci. 2008;87: 2329–2334. doi:10.3382/ps.2008-00193

20. Owens B, Tucker L, Collins MA, McCracken KJ. Effects of different feed additives alone or in combination on broiler performance, gut microflora and ileal histology. Br Poult Sci. 2008;49: 202–212. doi:10.1080/00071660802004890

21. Rodríguez ML, Rebolé A, Velasco S, Ortiz LT, Treviño J, Alzueta C. Wheat- and barley- based diets with or without additives influence broiler chicken performance, nutrient digestibility and intestinal microflora. J Sci Food Agric. 2012;92: 184–190. doi:10.1002/jsfa.4561

22. Classen HL, Campbell GL, Rossnagel BG, Bhatty R, Reichert RD. Studies on the use of hulless barley in chick diets: deleterious effects and methods of alleviation. Can J Anim Sci. 1985;65: 725–733. doi:10.4141/cjas85-085

23. McNab JM, Smithard RR. Barley β-Glucan: An antinutritional factor in poultry feeding. Nutr Res Rev. 1992;5: 45–60. doi:10.1079/NRR19920006

24. Ames N, Rhymer C, Rossnagel B, Therrien M, Ryland D, Dua S, et al. Utilization of diverse hulless barley properties to maximize food product quality. Cereal Food World. 2006; 51: 23–28. doi:10.1094/CFW-51-0023

25. Józefiak D, Kaczmarek S, Rutkowski A, Józefiak A, Jensen BB, Engberg RM. Fermentation in broiler chicken gastrointestinal tract as affected by high dietary inclusion of barley and by β-glucanase supplementation. J Anim Feed Sci. 2005;14: 695–704. doi:10.22358/jafs/67156/2005

26. Józefiak D, Rutkowski A, Kaczmarek S, Jensen BB, Engberg RM, Højberg O. Effect of β - glucanase and xylanase supplementation of barley- and rye-based diets on caecal microbiota of broiler chickens. Br Poult Sci. 2010;51: 546–557. doi:10.1080/00071668.2010.507243

27. Jiang Y, Zhang W, Gao F, Zhou G. Effect of sodium butyrate on intestinal inflammatory response to lipopolysaccharide in broiler chickens. Can J Anim Sci. 2015;95: 389–395. doi:10.4141/cjas-2014-183

28. Engberg RM, Hedemann MS, Leser TD, Jensen BB. Effect of zinc bacitracin and salinomycin on intestinal microflora and performance of broilers. Poult Sci. 2000;79: 1311– 1319. doi:10.1093/ps/79.9.1311

29. Dibner JJ, Richards JD. Antibiotic growth promoters in agriculture: history and mode of action. Poult Sci. 2005;84: 634–643. doi:10.1093/ps/84.4.634

30. Lee KW, Ho Hong Y, Lee SH, Jang SI, Park MS, Bautista DA, et al. Effects of anticoccidial and antibiotic growth promoter programs on broiler performance and immune status. Res Vet Sci. 2012;93: 721–728. doi:10.1016/j.rvsc.2012.01.001

31. Singh P, Karimi A, Devendra K, Waldroup PW, Cho KK, Kwon YM. Influence of penicillin on microbial diversity of the cecal microbiota in broiler chickens. Poult Sci. 2013;92: 272–276. doi:10.3382/ps.2012-02603

32. Olfert ED, Cross BM, McWilliam AA. Guide to the Care and Use of Experimental Animals. Vol. 1. 2nd ed. Canadian Council on Animal Care. Ottawa, ON, Canada; 1993.

33. CCAC Guidelines on: The Care and Use of Farm Animals in Research, Teaching and Testing. Canadian Council on Animal Care. Ottawa, ON, Canada.; 2009.

34. Ross 308 Broiler Performance Objectives. 2014. Available: http://en.aviagen.com/assets/Tech_Center/Ross_Broiler/Ross-308-Broiler-PO-2014-EN.pdf

35. Official methods of analysis, 18th. ed. Association of Official Analytical Chemists. Arlington, VA.; 2006.

36. Approved Methods of Analysis. 11th ed. American Association of Cereal Chemists, St. Paul, MN.; 2010.

37. ICC Standard Methods. International Association for Cereal Science and Technology, Vienna, Austria; 2011.

38. Boyd L, Holley R, Storsley J, Ames N. Effect of heat treatments on microbial load and associated changes to β-glucan physicochemical properties in whole grain barley. Cereal Chem. 2017;94: 333–340. doi:10.1094/CCHEM-04-16-0099-R

39. Zhao G, Nyman M, Jönsson JÅ. Rapid determination of short-chain fatty acids in colonic contents and faeces of humans and rats by acidified water-extraction and direct-injection gas chromatography. Biomed Chromatogr. 2006;20: 674–682. doi:10.1002/bmc.580

40. Osho SO, Wang T, Horn NL, Adeola O. Comparison of goblet cell staining methods in jejunal mucosa of turkey poults. Poult Sci. 2017;96: 556–559. doi:10.3382/ps/pew324

41. SAS User’s Guide: Statistics. Version 9.4 ed. SAS Institute. Inc., Cary, NC; 2008.

42. Holtekjølen AK, Vhile SG, Sahlstrøm S, Knutsen SH, Uhlen AK, Åssveen M, et al. Changes in relative molecular weight distribution of soluble barley beta-glucan during passage through the small intestine of pigs. Livest Sci. 2014;168: 102–108. doi:10.1016/j.livsci.2014.06.027

43. Gaborieau M, Castignolles P. Size-exclusion chromatography (SEC) of branched polymers and polysaccharides. Anal Bioanal Chem. 2011;399: 1413–1423. doi:10.1007/s00216-010-4221-7

44. Zhao Y, Zhou H-M, Huang Z-H, Zhao R-Y. Different aggregation states of barley β-glucan molecules affects their solution behavior: A comparative analysis. Food Hydrocoll. 2020;101: 105543. doi:10.1016/j.foodhyd.2019.105543

45. Lei F, Yin Y, Wang Y, Deng B, Yu HD, Li L, et al. Higher-level production of volatile fatty acids in vitro by chicken gut microbiotas than by human gut microbiotas as determined by functional analyses. Appl Environ Microbiol. 2012;78: 5763–5772. doi:10.1128/AEM.00327-12

46. Bautil A, Verspreet J, Buyse J, Goos P, Bedford MR, Courtin CM. Age-related arabinoxylan hydrolysis and fermentation in the gastrointestinal tract of broilers fed wheat-based diets. Poult Sci. 2019;98: 4606–4621. doi:10.3382/ps/pez159

47. Fry RE, Allred JB, Jensen LS, McGinnis J. Influence of enzyme supplementation and water treatment on the nutritional value of different grains for poults. Poult Sci. 1958;37: 372– 375. doi:10.3382/ps.0370372

48. Stanley D, Geier MS, Denman SE, Haring VR, Crowley TM, Hughes RJ, et al. Identification of chicken intestinal microbiota correlated with the efficiency of energy extraction from feed. Vet Microbiol. 2013;164: 85–92. doi:10.1016/j.vetmic.2013.01.030

49. Karunaratne ND, Ames NP, Van Kessel AG, Bedford MR, Newkirk RW, Classen HL. Effects of graded levels of hulless barley and beta-glucanase on beta-glucan depolymerization, digestive tract morphology and physiology of broiler chickens vaccinated for coccidiosis. Poult Sci. 2019;98(E-suppl. 1): 38.

50. Torok VA, Allison GE, Percy NJ, Ophel-Keller K, Hughes RJ. Influence of antimicrobial feed additives on broiler commensal posthatch gut microbiota development and performance. Appl Environ Microbiol. 2011;77: 3380–3390. doi:10.1128/AEM.02300-10

51. Simon K, Verwoolde MB, Zhang J, Smidt H, de Vries Reilingh G, Kemp B, et al. Long-term effects of early life microbiota disturbance on adaptive immunity in laying hens. Poult Sci. 2016;95: 1543–1554. doi:10.3382/ps/pew088

52. Xiong W, Wang Y, Sun Y, Ma L, Zeng Q, Jiang X, et al. Antibiotic-mediated changes in the fecal microbiome of broiler chickens define the incidence of antibiotic resistance genes. Microbiome. 2018;6. doi:10.1186/s40168-018-0419-2

53. Beckmann L, Simon O, Vahjen W. Isolation and identification of mixed linked beta-glucan degrading bacteria in the intestine of broiler chickens and partial characterization of respective 1,3-1,4-beta-glucanase activities. J Basic Microbiol. 2006;46: 175–185. doi:10.1002/jobm.200510107

54. Pieper R, Bindelle J, Malik G, Marshall J, Rossnagel BG, Leterme P, et al. Influence of different carbohydrate composition in barley varieties on *Salmonella* Typhimurium var. Copenhagen colonisation in a “Trojan” challenge model in pigs. Arch Anim Nutr. 2012;66: 163–179. doi:10.1080/1745039X.2012.676814

55. Gorham JB, Kang S, Williams BA, Grant LJ, McSweeney CS, Gidley MJ, et al. Addition of arabinoxylan and mixed linkage glucans in porcine diets affects the large intestinal bacterial populations. Eur J Nutr. 2017;56: 2193–2206. doi:10.1007/s00394-016-1263-4

56. Boros D, Marquardt RR, Slominski BA, Guenter W. Extract viscosity as an indirect assay for water-soluble pentosan content in rye. Cereal Chem. 1993 [cited 13 May 2020]. Available: https://agris.fao.org/agris-search/search.do?recordID=US9433814

57. Saulnier L, Peneau N, Thibault J-F. Variability in grain extract viscosity and water-soluble arabinoxylan content in wheat. J Cereal Sci. 1995;22: 259–264. doi:10.1006/jcrs.1995.0062

58. Knudsen KEB. The impact of arabinoxylan and β-glucan in the nutrition of pigs and poultry. Proc 37th West Nutr Conf. 2016; 130–142.

59. Morel PCH, Melai J, Eady SL, Coles GD. Effect of non-starch polysaccharides and resistant starch on mucin secretion and endogenous amino acid losses in pigs. Asian Australas J Anim Sci. 2005;18: 1634–1641. doi:2005.18.11.1634

60. Cadogan DJ, Choct M. Pattern of non-starch polysaccharide digestion along the gut of the pig: Contribution to available energy. Anim Nutr. 2015;1: 160–165. doi:10.1016/j.aninu.2015.08.011

61. Mälkki Y, Virtanen E. Gastrointestinal effects of oat bran and oat gum: a review. LWT-Food Sci Technol. 2001;34: 337–347. doi:10.1006/fstl.2001.0795

62. Karunaratne ND, Bedford MR, Newkirk RW, Classen HL. Graded levels of hulless barley and β-glucanase affect the digestive tract morphology of broiler chickens vaccinated for coccidiosis. Poult Sci. 2017a;96 (E-suppl. 1): 109.

63. Brenes A, Smith M, Guenter W, Marquardt RR. Effect of enzyme supplementation on the performance and digestive tract size of broiler chickens fed wheat- and barley-based diets. Poult Sci. 1993;72: 1731–1739. doi:10.3382/ps.0721731

64. Jørgensen H, Zhao XQ, Knudsen KE, Eggum BO. The influence of dietary fibre source and level on the development of the gastrointestinal tract, digestibility and energy metabolism in broiler chickens. Br J Nutr. 1996;75: 379–395. doi:10.1079/bjn19960141

65. Salih ME, Classen HL, Campbell GL. Response of chickens fed on hull-less barley to dietary β-glucanase at different ages. Anim Feed Sci Technol. 1991;33: 139–149. doi:10.1016/0377-8401(91)90052-T

66. Almirall M, Esteve-Garcia E. Rate of passage of barley diets with chromium oxide: influence of age and poultry strain and effect of beta-glucanase supplementation. Poult Sci. 1994;73: 1433–1440. doi:10.3382/ps.0731433

67. Hesselman K, Åman P. The effect of β-glucanase on the utilization of starch and nitrogen by broiler chickens fed on barley of low- or high-viscosity. Anim Feed Sci Technol. 1986;15: 83–93. doi:10.1016/0377-8401(86)90015-5

68. Ravindran V, Tilman ZV, Morel PCH, Ravindran G, Coles GD. Influence of β-glucanase supplementation on the metabolisable energy and ileal nutrient digestibility of normal starch and waxy barleys for broiler chickens. Anim Feed Sci Technol. 2007;134: 45–55. doi:10.1016/j.anifeedsci.2006.04.012

69. Miles RD, Butcher GD, Henry PR, Littell RC. Effect of antibiotic growth promoters on broiler performance, intestinal growth parameters, and quantitative morphology. Poult Sci. 2006;85: 476–485. doi:10.1093/ps/85.3.476

70. Selber-Hnatiw S, Rukundo B, Ahmadi M, Akoubi H, Al-Bizri H, Aliu AF, et al. Human gut microbiota: toward an ecology of disease. Front Microbiol. 2017;8. doi:10.3389/fmicb.2017.01265

71. Donaldson GP, Lee Sm, Mazmanian SK. Gut biogeography of the bacterial microbiota. Nat Rev Microbiol. 2016;14: 20–32. doi:http://dx.doi.org/10.1038/nrmicro3552

72. Croom J, Chichlowski M, Froetschel M, McBride BW, Qui R, Koci MD. The effects of direct-fed microbial, primalac, or salinomycin supplementation on intestinal lactate isomers and cecal volatile fatty acid concentrations in broilers. Int J Poult Sci. 2009;8: 128–132.

73. Smirnov A, Perez R, Amit-Romach E, Sklan D, Uni Z. Mucin dynamics and microbial populations in chicken small intestine are changed by dietary probiotic and antibiotic growth promoter supplementation. J Nutr. 2005;135: 187–192. doi:10.1093/jn/135.2.187

74. Danzeisen JL, Kim HB, Isaacson RE, Tu ZJ, Johnson TJ. Modulations of the chicken cecal microbiome and metagenome in response to anticoccidial and growth promoter treatment. Parkinson J, editor. PLoS one. 2011;6: e27949. doi:10.1371/journal.pone.0027949

75. Cardoso V, Ferreira AP, Costa M, Ponte PIP, Falcão L, Freire JP, et al. Temporal restriction of enzyme supplementation in barley-based diets has no effect in broiler performance. Anim Feed Sci Technol. 2014;198: 186–195. doi:10.1016/j.anifeedsci.2014.09.007

76. Rada V, Marounek M. Effect of monensin on the crop microflora of broiler chickens. Ann Zootech. 1996;45: 283–288.

77. Caspary WF. Physiology and pathophysiology of intestinal absorption. Am J Clin Nutr. 1992;55: 299S–308S. doi:10.1093/ajcn/55.1.299s

78. Sukhotnik I, Coran AG, Mogilner JG, Shamian B, Karry R, Lieber M, et al. Leptin affects intestinal epithelial cell turnover in correlation with leptin receptor expression along the villus-crypt axis after massive small bowel resection in a rat. Pediatr. Res. 2009;66: 648– 653. doi:10.1203/PDR.0b013e3181be9f84

79. Seno H, Miyoshi H, Brown SL, Geske MJ, Colonna M, Stappenbeck TS. Efficient colonic mucosal wound repair requires Trem2 signaling. Proc Natl Acad Sci USA. 2009;106: 256– 261. doi:10.1073/pnas.0803343106

80. Kuhn KA, Manieri NA, Liu T-C, Stappenbeck TS. IL-6 stimulates intestinal epithelial proliferation and repair after injury. PLoS one. 2014;9: e114195. doi:10.1371/journal.pone.0114195

81. Bamford KB. Chronic gastrointestinal inflammation. FEMS Immunol Med Microbiol. 1999;24: 161–168. doi:10.1111/j.1574-695X.1999.tb01277.x

82. Ward PA, Lentsch AB. The acute inflammatory response and its regulation. Arch Surg. 1999;134: 666–669. doi:10.1001/archsurg.134.6.666

83. Hu Z, Guo Y. Effects of dietary sodium butyrate supplementation on the intestinal morphological structure, absorptive function and gut flora in chickens. Anim Feed Sci Technol. 2007;132: 240–249. doi:10.1016/j.anifeedsci.2006.03.017

84. Kim MH, Kang SG, Park JH, Yanagisawa M, Kim CH. Short-chain fatty acids activate GPR41 and GPR43 on intestinal epithelial cells to promote inflammatory responses in mice. Gastroenterology. 2013;145: 396–406.e1–10. doi:10.1053/j.gastro.2013.04.056

85. Macia L, Tan J, Vieira AT, Leach K, Stanley D, Luong S, et al. Metabolite-sensing receptors GPR43 and GPR109A facilitate dietary fibre-induced gut homeostasis through regulation of the inflammasome. Nat Commun. 2015;6: 6734. doi:10.1038/ncomms7734

86. Hume ME, Clemente-Hernández S, Oviedo-Rondón EO. Effects of feed additives and mixed *Eimeria* species infection on intestinal microbial ecology of broilers. Poult Sci. 2006;85: 2106–2111. doi:10.1093/ps/85.12.2106

87. Macdonald SE, Nolan MJ, Harman K, Boulton K, Hume DA, Tomley FM, et al. Effects of *Eimeria tenella* infection on chicken caecal microbiome diversity, exploring variation associated with severity of pathology. PLoS one. 2017;12: e0184890. doi:10.1371/journal.pone.0184890

88. Leung H, Yitbarek A, Snyder R, Patterson R, Barta JR, Karrow N, et al. Responses of broiler chickens to *Eimeria* challenge when fed a nucleotide-rich yeast extract. Poult Sci. 2019;98: 1622–1633. doi:10.3382/ps/pey533

89. Apajalahti J. Comparative gut microflora, metabolic challenges, and potential opportunities. J Appl Poult Res. 2005;14: 444–453. doi:10.1093/japr/14.2.444

90. Radu J, Van Dijk C, Wheelhouse RK, Hummant CA, Gadbois P. Feed and water consumption and performance of male and female broilers fed salinomycin and maduramicin followed by a withdrawal ration. Poult Sci. 1987;66: 1878–1881. doi:10.3382/ps.0661878

91. Elwinger K, Berndtson E, Engström B, Fossum O, Waldenstedt L. Effect of antibiotic growth promoters and anticoccidials on growth of *Clostridium perfringens* in the caeca and on performance of broiler chickens. Acta Vet Scand. 1998;39: 433–441.

92. Dumonceaux TJ, Hill JE, Hemmingsen SM, Van Kessel AG. Characterization of intestinal microbiota and response to dietary virginiamycin supplementation in the broiler chicken. Appl Environ Microbiol. 2006;72: 2815–2823. doi:10.1128/AEM.72.4.2815-2823.2006

93. van Der Wielen PW, Biesterveld S, Notermans S, Hofstra H, Urlings BA, van Knapen F. Role of volatile fatty acids in development of the cecal microflora in broiler chickens during growth. Appl Environ Microbiol. 2000;66: 2536–2540. doi:10.1128/aem.66.6.2536-2540.2000

94. Pourabedin M, Zhao X. Prebiotics and gut microbiota in chickens. FEMS Microbiol Lett. 2015;362. doi:10.1093/femsle/fnv122

95. Panda AK, Rao SVR, Raju MVLN, Sunder GS. Effect of butyric acid on performance, gastrointestinal tract health and carcass characteristics in broiler chickens. Asian Australas J Anim Sci. 2009;22: 1026–1031. doi:10.5713/ajas.2009.80298

96. Wu W, Xiao Z, An W, Dong Y, Zhang B. Dietary sodium butyrate improves intestinal development and function by modulating the microbial community in broilers. PLoS one. 2018;13. doi:10.1371/journal.pone.0197762

97. Karunaratne ND, Bedford MR, Newkirk RW, Classen HL. Graded levels of hulless barley and β-glucanase affect the performance of broiler chickens vaccinated for coccidiosis in an age dependent manner. Poult Sci. 2017b;96 (E-suppl. 1): 50.

